# Mapping cancer gene dynamics through state-specific interactions

**DOI:** 10.1101/2025.01.08.631858

**Authors:** Adrián Maqueda-Real, Laia Ollé-Monràs, Solip Park

## Abstract

Metastatic cancer, a major cause of mortality, has been understudied compared to primary tumors, leaving gaps in our understanding of how cancer genes adapt between these states. We analyzed the association between mutations and copy-number alterations in 25,000 tumor samples from both primary and metastatic cancers. Our findings show that cancer genes display distinct interaction strengths across these states, with 27.45% of genes, including *ARID1A*, *FBXW7*, and *SMARCA4*, shifting between one-hit and two-hit drivers. Interaction strengths varied by cancer state and treatment conditions, revealing seven state-specific interactions. We also identified 38 primary-specific and 21 metastatic-specific high-order interactions, enriched in cancer hallmarks, indicating unique tumor progression mechanisms. These findings highlight dynamic tumor progression mechanisms and underscore the importance of considering cancer state in research and treatment strategies for precise therapeutic interventions.

## Introduction

Cancer is one the most prevalent diseases, affecting one in every six individuals worldwide. To gain a comprehensive understanding of cancer, substantial efforts have been thoroughly pursued, leveraging rapidly advancing sequencing technologies. These efforts include collaborative initiatives such as The Cancer Genome Atlas (TCGA), the Pan-Cancer Analysis of Whole Genomes (PCAWG), along with contributions like Memorial Sloan Kettering – Metastatic Events and Tropisms (MSK-MET) and Hartwig Medical Foundation (Consortium, 2020; Ding et al., 2018; Nguyen et al., 2022; Priestley et al., 2019). The volume of sequencing data for cancer patients continues to grow, with Integrative OncoGenomics (IntOGen) database (Martinez-Jimenez et al., 2020) reporting over 33,000 samples from 266 cohorts as of May 2023.

Despite this extensive collection of data and numerous studies aiming to uncover the characteristics of tumors across various cancer types, there is a significant bias in most studies, predominantly focusing on primary tumors (comprising 84.1% of collected samples in IntOGen) and untreated conditions (84.6%). This focus is paradoxical given that more than 90% of cancer-related deaths are attributed to metastatic cancers (Lambert et al., 2017). This bias leaves the true disparities between primary and metastatic tumors largely unexplored.

In recent years, several extensive studies have sought to elucidate the genomic differences between primary and metastatic cancers to understand their unique characteristics (Consortium, 2017; Martinez-Jimenez et al., 2023; Nguyen et al., 2022). Large-scale comparisons between primary and metastatic cancer have revealed distinct characteristics of metastatic tumors, including higher mutational burden, increased genomic instability, and occurrences of whole-genome duplication (WGD), which are more prevalent compared to primary tumors (Martinez-Jimenez et al., 2023; Nguyen et al., 2022). For instance, the MSK-MET study assembled a pan-cancer cohort comprising over 25,000 samples (Nguyen et al., 2022). This diverse dataset, encompassing both primary (15,632 samples) and metastatic samples (10,143 samples), has provided valuable insights into variations in chromosomal instability and driver alterations relevant to tumor progression. Another significant undertaking involved whole-genome sequencing of 5,365 samples from 23 cancer types, spanning unpaired primary samples (1,914 samples from the PCAWG) and metastatic samples (3,451 samples from the Hartwig Medical Foundation)(Martinez-Jimenez et al., 2023). The objective of this effort was to comprehensively characterize the genomic distinctions between the two cancer states in a cancer-type specific manner and under the influence of exposure to cancer treatments. In parallel, specific cancer-focused studies have explored genomic diversity by comparing primary and metastatic tumors. These studies frequently employ phylogenetic tree reconstructions to elucidate the trajectory of tumor progression (Bertucci et al., 2019; Brastianos et al., 2015; Brown et al., 2017; Eckert et al., 2016; Hu et al., 2020; Jimenez-Sanchez et al., 2017; Makohon-Moore et al., 2017; Naxerova et al., 2017; Noorani et al., 2020; Reiter et al., 2020; Robinson et al., 2017; Shih et al., 2020).

While these studies made substantial progress in understanding the genomic differences of cancer states, there remains a gap in research that quantitatively examines how cancer fitness levels may change in relation to specific cancer states across cancer driver genes or cancer types. This extends beyond the examination of individual genomic alterations unique to a particular cancer state. Importantly, through an extensive analysis of > 10,000 human cancer exomes, our recent study demonstrated that cancer genes have the capacity to adapt their optimal cancer fitness level varying on their cellular contexts, such as cancer types (Park et al., 2021). Moreover, we proposed that high-order genetic interactions — specifically, two-way interactions between mutations and copy-number changes within a gene, which perturb depending on the mutation status of another gene — play an important role in switching the mode of action of cancer genes (either two-hit or one-hit driver). Collectively, these findings provide a comprehensive explanation for the diverse variability observed in tumorigenesis.

To explore how cancer genes change their mode of action depending on cancer states, such as primary and metastatic tumors, we systematically analyzed their preferences for different genomic alterations within a cancer gene across various cancer types using the MSK-MET dataset (Nguyen et al., 2022). This dataset is not only well-balanced between primary and metastatic tumors but is also the largest cohort available to date. By quantifying the strength of two-way interactions, we identified state-specific cancer types that exhibit stronger fitness levels in either primary or metastatic states. Furthermore, we demonstrated that a significant portion of cancer genes (27.45% of tested genes) alter their mode of action depending on the cancer state to achieve optimal fitness levels. This includes *ARID1A*, *FBXW7*, and *SMARCA4*, which are key to understanding the progression from primary to metastatic tumors. Additionally, we identified cancer-state-specific high-order interactions that modulate the strength of two-way interactions through other genes to achieve state-specific fitness levels. Remarkably, we established the clinical impact of various associations among different genomic alterations within a gene or between genes by analyzing the prognosis of cancer patients. Our findings suggest that this approach provides a powerful model for understanding the distinct characteristics of cancer states and advancing knowledge of the condition-specific cancer fitness landscape. Our cancer-state specific analysis can be interactively visualized at https://primet-fitness.bioinfo.cnio.es/.

## Results

### Cancer state-specific changes in interactions between mutations and CNAs

To quantify the cancer fitness level in each cancer statement (i.e., primary and metastasis), we examined the co-occurrence of mutations (specifically actionable mutations identified by OncoKB (Chakravarty et al., 2017); see **Methods**) and copy-number alterations (CNAs, either loss or gain) for each gene in cancer types in which these alterations were frequent (**Figure 1A**). We employed a consistent approach to analyze genomic data from 25,420 tumors across 10 different cancer types including their sub-types (*N*=50; **Tables S1**). These data are part of the Memorial Sloan Kettering - Metastatic Events and Tropisms (MSK-MET) project (Nguyen et al., 2022), consisting of 15,387 primary samples (60.53%) and 10,033 metastatic samples (39.47%; **Figure 1B**). We applied the log-linear regression model for measuring the interactions between mutations and CNAs within a gene (i.e., two-way interaction), as described in our previous study (Park et al., 2021), to 384 cancer driver genes to investigate changes in cancer fitness between primary and metastatic tumors (**Figure 1A**; **Tables S2**). These genes include 195 tumor suppressor genes (TSGs), 167 oncogenes (OGs), and 22 dual-functional genes (DFGs) (see **Methods**). Specifically, we tested genes that had a mutation frequency of > 1% and a CNA frequency of > 10% within a single cancer type, considering the specific cancer state. (See, **Methods**). On average, genes were tested for two-way interaction in 2.33 cancer types for primary tumors and 2.68 for metastatic tumors. In total, 148 genes were examined (126 in primary and 120 in metastatic tumors, with 90 genes overlapping) across 650 gene-cancer type pairs (299 in primary and 351 in metastatic tumors). Initially, we investigated the overall two-way interaction strengths (effect sizes) between mutations and CNAs within genes, focusing on differences between primary and metastatic tumors across ten cancer types.

**Figure 1.**
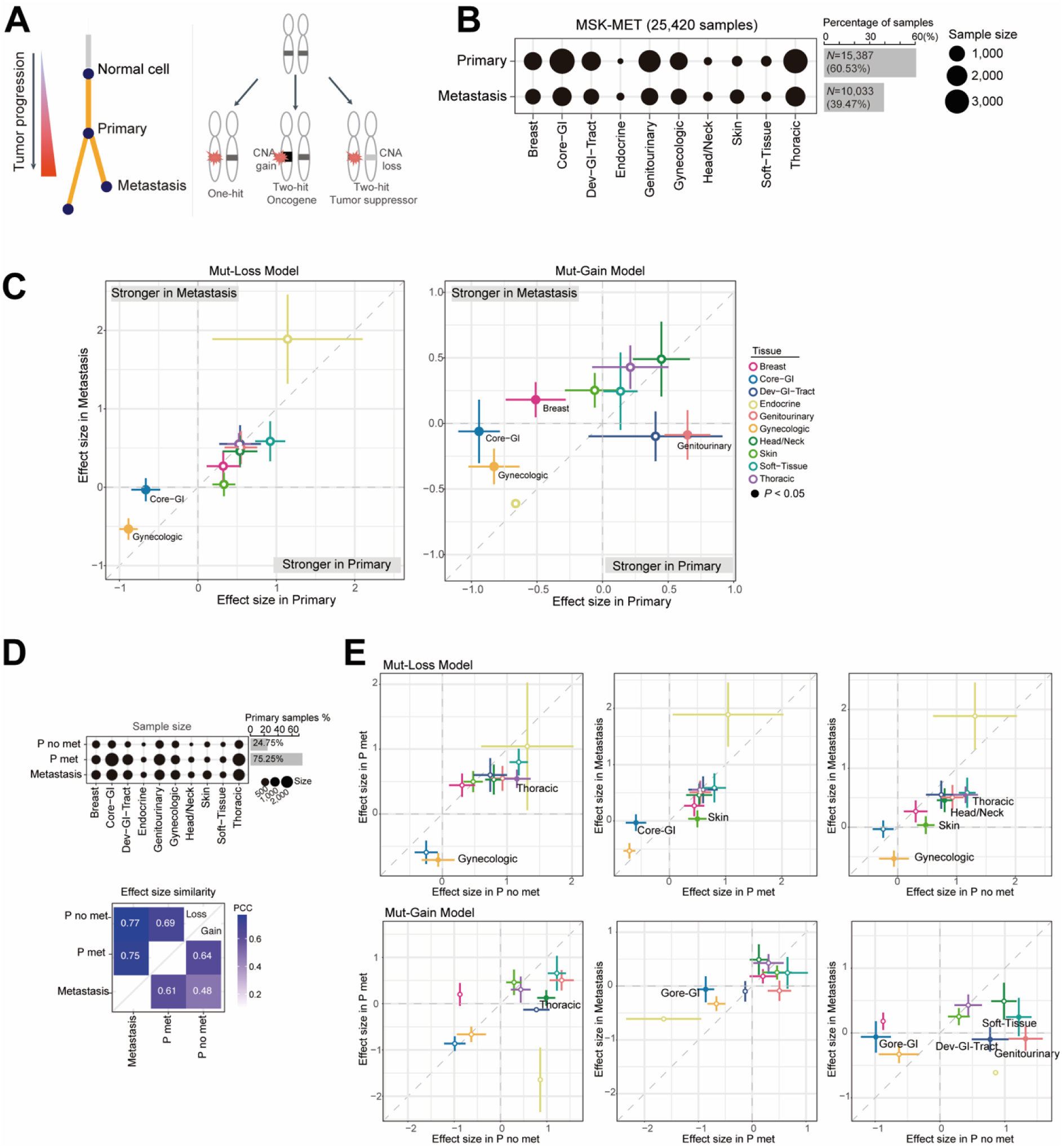
Interactions between mutations and CNAs within a gene across cancer types in both primary and metastatic tumors. (**a**) Three possible cancer fitness models for the association between mutation and CNAs within a cancer gene. (**b**) An overview of the sample size distribution across cancer types. (**c**) The average interaction strength between mutation and CNAs within a gene in primary (x-axis) and metastatic tumors (y-axis). Error bars indicate the standard error for primary and metastatic tumors. Cancer types with statistically significant differences in interaction strength between primary and metastatic tumors are denoted with filled circles by Mann–Whitney *U*-test (*P*-val < 0.05). (**d**) The size distribution of tested samples across conditions: primary samples from non-metastatic patients (P no met), primary tumors from metastatic patients (P met), and metastatic samples. The heat map displays the similarities of interaction strength between different cancer states in loss and gain model, separately. (**e**) Average interaction strength between mutations and CNAs within a gene across different cancer types in two distinct cancer states. Tumor types with statistically significant differences between two cancer states are indicated with filled circles using Mann–Whitney *U*-test (*P*-val < 0.05).

Metastatic tumors generally exhibited a higher overall two-way interaction strength than primary tumors, except for Genitourinary cancer, which showed an average effect size of 0.64 in primary tumors versus -0.09 in metastatic tumors (*P* = 4.40E-2 by Mann–Whitney *U*-test) in the CNA-gain model. Specifically, three cancer types (3/10; Breast, Core-GI, and Gynecologic cancer) displayed significantly higher interaction strengths in metastatic tumors in at least one of the CNA models (*P* < 0.05 by Mann–Whitney U-test; **Figure 1C** and **Figure S1**). This includes, in the CNA-loss model, Core-GI (-0.66 versus -0.03; *P* = 6.07E-3) and Gynecologic cancer (-0.89 versus -0.53; *P* = 1.74E-2), and in the CNA-gain model, Breast (-0.51 versus 0.18; *P* = 3.02E-2), Core-GI (-0.94 versus -0.06; *P* = 1.46E-3), and Gynecologic cancer (-0.83 versus -0.33; *P* = 2.56E-2).

Importantly, the interaction strengths were not significantly influenced by potential confounding factors, such as patient age, sample purity, sequencing coverage, and sample clonality (**Figure S2A-D**). Additionally, specific genomic alterations, which differed significantly between primary and metastatic samples from the original MSK-MET study (Nguyen et al., 2022), did not account for the observed variation in interaction strengths (**Figure S2E**). This suggests that the differences observed in cancer states in our analysis are principally driven by the diversity of various cancer genes examined and the combination of two distinct genomic events, rather than by specific genes or individual genomic alterations (either mutation or CNAs only).

These differences in interaction strengths between primary and metastatic tumors were also evident in matched subtypes (**Figure S2 F** and **G**). In the CNA-loss model, significant differences were observed in Gynecologic clear cell ovarian cancer (CCOV, *P* = 1.02E-2) and in the CNA-gain model as well, including Core-GI-Colorectal MSS (CRC-MSS, *P* = 3.03E-3), Core-GI-Small Bowel cancer (SBS, *P* = 1.47E-2), and Gynecologic UCEC-Serous (UCEC-SEROUS, *P* = 4.50E-3). Interestingly, several cancer subtypes exhibited significantly different interaction strengths between primary and metastatic tumors, distinctions that were not evident at the cancer type level. The subtypes identified in the CNA-loss model include Breast-Duct-TN (IDC-TN, *P* = 2.97E-2), Developmental GI Tract-Cholangio Extrahepatic (CHOL-extra, *P* = 6.35E-3), Developmental GI Tract-Gallbladder Cancer (GBC, *P* = 2.03E-2), Skin-Mucosal (MUCM, *P* = 3.45E-2), and Melanoma Cutaneous (SKCM, *P* = 3.37E-3). Meanwhile, the CNA-gain model revealed subtypes such as Skin-Cutaneous Squamous Cell (CSCC, *P* = 2.16E-3), Genitourinary Upper Tract Urothelial Carcinoma (UTUC, *P* = 1.52E-4), and Endocrine-Thyroid Poorly Differentiated (THPD, *P* = 3.11E-2).

To further explore the differences between primary and metastatic tumors, we divided primary tumors into two groups based on their origins: tumors from non-metastatic patients (P no met) and those from metastatic patients (P met) to comprehensively understand the changes in interaction strengths across various cancer types. This categorization indicated that 24.75% of primary samples were from non-metastatic patients (ranging from 13.61% to 47.15% across cancer types), while 75.25% were from metastatic patients (ranging from 52.85% to 86.39% across cancer types). As expected, metastatic tumors exhibited a higher similarity to primary tumors from metastatic patients (P met; PCC = 0.75 in the loss model, 0.61 in the gain model) compared to those from non-metastatic patients (P no met; PCC = 0.77 in the loss model, 0.48 in the gain model). Categorizing primary tumors based on their metastatic or non-metastatic status revealed an increased number of distinct cancer types. Eight of these cancer types (80%) showed notable differences in at least one of the loss or gain models when comparing primary tumors from non-metastatic patients (P no met) and metastatic patients. This contrasts with only two cancer types showing significant differences when comparing primary tumors from metastatic patients (P met) versus metastatic patients, or primary tumors from non-metastatic patients (P no met) versus primary tumors from metastatic patients (P met) conditions (*P* < 0.05 by the Mann–Whitney *U*-test, **Figure 1E**). It underscores the critical role of accurate classification in discerning interaction strength between primary and metastatic tumors, highlighting their potential importance in understanding tumor evolution.

### Perturbation of gene classes in primary and metastatic tumors

We subsequently directed our focus towards elucidating differences in interaction strength between mutations and CNAs within cancer-related genes, specifically comparing primary and metastatic tumors across ten cancer types (**Figure 2A**). In total, we detected significant two-way interactions in 27 genes, involving 51 pairs across cancer types in primary tumors (21.43% of tested genes and 17.06% of tested pairs) and 42 genes in 73 pairs in metastatic tumors (35% of tested genes and 20.80% of tested pairs) at a permutation-based false discovery rate (FDR < 10%, **Figure S3** and **Tables S2**). Additionally, 34 interactions in 20 genes overlapped between primary and metastatic tumors (31 in the CNA-Loss model and 3 in the CNA-Gain model). At the current sample size, the saturation analysis suggests that the number of interactions in both primary and metastatic tumors has not reached saturation (**Figure S4A**), indicating the need for more samples to achieve a complete understanding of two-way interactions in both primary and metastatic tumors.

**Figure 2.**
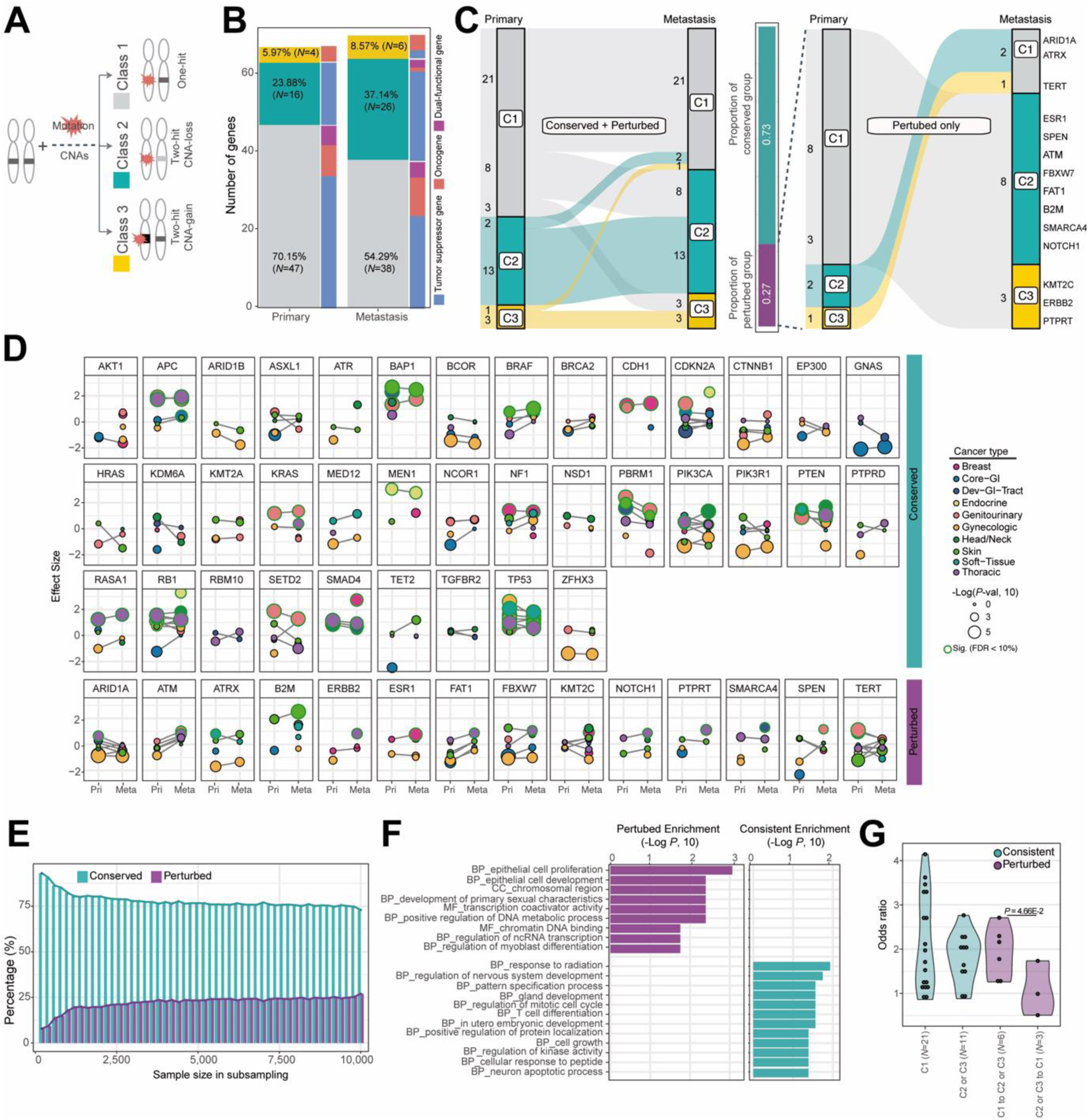
Perturbation of two-way interactions between primary and metastatic tumors. (**a**) Gene classification based on their preferences for two-way interactions across tested cancer types. (**b**) The number of genes assigned to each gene class in primary and metastatic tumors. (**c**) The number of genes categorized as consistent (maintaining their preferences for two-way interactions between primary and metastatic tumors) and perturbed (changing their preferences for two-way interaction). (**d**) The distribution of interaction strength across tested cancer types between primary and metastatic tumors in the consistent group (top panel) and the perturbed group (bottom panel). A higher interaction strength indicates stronger co-occurrence between mutations and CNAs within a gene. (**e**) The average percentage of genes in the conserved and perturbed groups as determined by subsampling analysis. Samples were randomly selected, increasing by 200 samples each time until reaching the final sample size (100 iterations). Error bars indicate the standard error. (**f**) Significantly enriched GO terms in the consistent versus perturbed groups (Fisher’s one-tailed test, *P* < 0.05). Three categories of GO terms were considered: biological process (BP), cellular component (CC) and molecular function (MF). (**g**) Odds ratio of clonal versus sub-clonal variants in two-way interactions between primary and metastatic tumors. A higher odds ratio indicates a greater prevalence of clonal variants in metastatic tumors compared to primary tumors. Mann–Whitney *U*-test result is provided for odds ratio comparisons between groups.

To investigate the details of these interaction changes across genes between different cancer states, we categorized genes into three classes based on their preference for interactions between mutations and CNAs, separately for primary and metastatic states. These classes align with the definitions proposed by our previous study (Park et al., 2021) at an FDR 10%: (i) ‘One-Hit Driver (genes that do not exhibit significant co-occurrence between mutation and CNAs in any of the tested cancer types, Class 1), (ii) ‘Two-Hit Loss’ (genes that only show significant interaction between mutation and CNA-loss in at least one tested cancer types, Class 2), and (iii) ‘Two-Hit Gain’ (genes that only show significant interaction between mutation and CNA-gain in at least one tested cancer type, Class 3). In this study, we did not observe any genes with significant interactions between mutations and both CNA-loss and CNA-gain, either within the same cancer type or across different cancer types. We considered genes tested in at least two cancer types: 67 genes in primary and 70 genes in metastatic tumors, with 51 genes overlapping in both conditions (**Tables S3**). The distribution of each class is comparable between the two cancer states (**Figure 2B**). In the primary state, 70.15% of genes are assigned to Class 1, 23.88% to Class 2, and 5.97% to Class 3. Similarly, in the metastatic state, 54.29% of genes are assigned to Class 1, 37.14% to Class 2, and 8.57% to Class 3.

In accordance with the two-hit model of cancer genes, we observed that TSGs were predominantly enriched in Class 2 (100% in primary and 88.46% in metastatic tumors), which includes classical tumor suppressor genes like *TP53*, *APC*, *PTEN*, *RB1*, and *NF1*. Many oncogenes were also categorized into Class 3 (100% in primary and 66.67% in metastatic tumors), including *KRAS*, *BRAF*, and *PIK3CA*. Furthermore, we verified that mutated alleles are commonly amplified in Class 3 genes in both cancer states (80.0% of mut-CNA gain interactions from Mann–Whitney *U*-test test with a *P*-value < 0.05; **Figure 5**). This observation aligns with previous findings regarding the positive selection of mutant allele imbalance in oncogenic variants (Bielski et al., 2018). Notably, we observed a strong enrichment of essential genes in Class 1 in both primary (12.8%) and metastatic tumors (15.8%; **Figure S4C**), following the frequencies of Class 2 (12.5% in primary and 7.69% in metastatic tumors). Interestingly, the enrichment of essential genes in Class 1 was more pronounced in metastatic than primary tumors, with marginal significance (odds ratio = 2.44, Fisher’s exact test *P*-value = 0.19). This suggests that essential genes shift to Class 1 from primary to metastatic tumors, indicating changes in their mode of action for adapting optimal activity fitness level depending on the cancer state

In our exploration of changes in interactions depending on the cancer state, we found that many of the compared cancer genes which were tested both states retained their classifications between cancer states (72.55%, 37 out of 51; referred to as the ’consistent group’), with 56.76% in Class 1, 35.14% in Class 2, and 8.11% in Class 3. However, a substantial portion exhibited changes in class between primary and metastatic tumors (27.45%, 14 out of 51; referred to as the ’perturbed group’, **Figure 2C** and **D**). Notably, in a saturation analysis, the distribution of gene classes and the proportions of the consistent and perturbed groups are maintained, even though the number of significant interactions has not reached saturation with the current sample size (**Figure 2E**; **Figure S4A** and **B**). Moreover, this trend is remained consistent the same to the analysis after separating primary samples into two groups (from metastatic patients and from non-metastatic patients; **Figure 4D** and **E**), suggesting that the observed proportion of perturbed genes is stable regardless of the sample size or the number of significant interactions (**Tables S4** and **5**).

**Figure 4.**
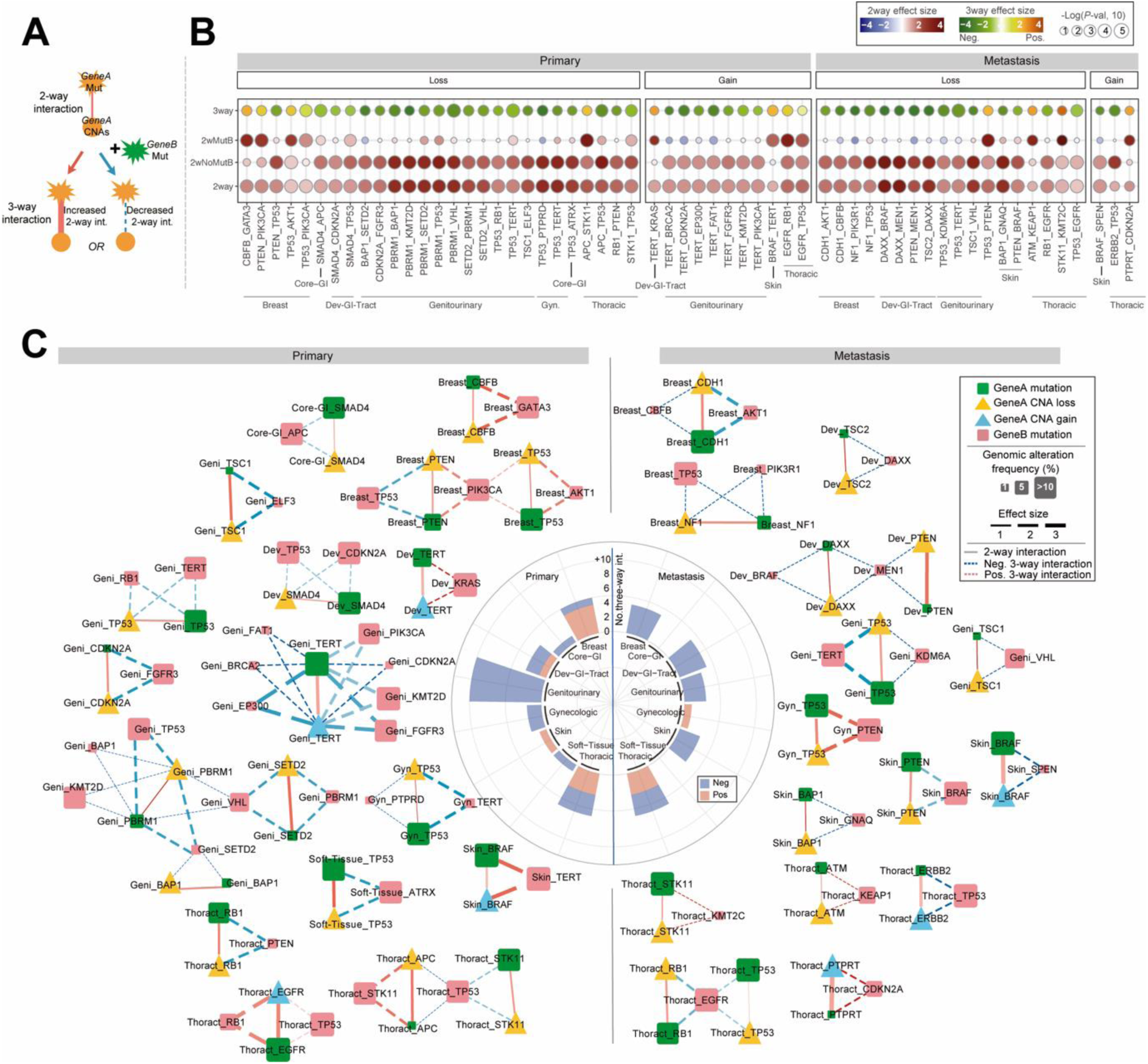
Cancer-state specific third-order genetic interactions across cancer types. (**a**) A third-order genetic interaction refers to a situation where the strength of a two-way interaction involving the first gene changes based on the mutation status of a second gene, as determined using a log-linear regression model. (**b**) Interaction strength and *P*-values for the 59 identified third-order interactions. These interactions include: the relationship between mutation and CNAs of the first gene depending on the mutation status of a second gene (third-order interactions), interaction between mutation and CNAs of the first gene with second gene mutations (2wMutB), without second gene mutations (2wNoMutB), and considering all samples without regard to the second gene’s mutation status (2way). (**c**) Third-order interaction networks between primary and metastatic tumors across various cancer types. Nodes represent the type of genomic events (squares for mutations and triangles for CNAs), with sizes indicating the frequency of alterations in tested cancer types and colors representing the type of CNAs (yellow for CNA loss and blue for CNA gain) or the type of associated genes (green for the first gene and red for the second gene). Edge types indicate the interaction type (solid line for two-way interactions and dashed line for three-way interactions). Edge colors represent the type of third-order interaction (blue for negative and red for positive), and edge widths indicate the strength of the interaction (absolute log of odds ratio). All networks were visualized using Cytoscape (Shannon et al., 2003).

Interestingly, the consistent and perturbed groups exhibit distinct Gene Ontology (GO) enrichments (**Figure 2F**). The perturbed group shows notable enrichment in processes related to epithelial cell proliferation, chromatin regulation, and other gene regulation processes. Conversely, cell cycle regulation, cell growth, and regulation of kinase activity are specifically enriched in the consistent group (*P* < 0.05 by one-sided Fisher’s exact test). These findings suggest that each group contributes uniquely to both common functional roles and cancer state-specific functional roles necessary for achieving the optimal cancer fitness level. Furthermore, we investigated whether the two groups (consistent versus perturbed) exhibited different clonalities across cancer states by calculating the odds ratio of clonal versus subclonal variants between primary and metastatic tumors. Surprisingly, both the Class 1 consistent group and the Class 2 or 3 consistent group showed similar odds ratios for clonal compared to sub-clonal variants from primary to metastatic state (median for Class 1 = 1.84, median for Class 2 or Class 3 = 1.98; *P* = 0.35 from a Mann–Whitney *U*-test). Additionally, the group transitioning from one-hit to two-hits (from Class 1 in primary to Class 2 or 3 in metastasis) displayed a relatively similar odds ratios to the consistent groups (median odds ratio = 1.97; *P*-value compared to the consistent group = 0.45 from a Mann–Whitney *U*-test). Notably, the perturbed group transitioning from two-hits to one-hit (from Class 2 or 3 to Class 1; including *ATRX*, *ARID1A* and *TERT*) exhibited significantly smaller odds ratios (median odds ratio = 0.99; *P*-value = 4.66E-2 from a Mann–Whitney *U*-test), suggesting that clonal variants were more prevalent in primary tumors compared to metastatic tumors (**Figure 2G**). This indicates that a large proportion of this group may consist of early mutations, resulting in perturbations from two-hits in primary tumors to one-hit in metastatic tumors. Our findings could align with previous studies, such as the three times increased frequency of early clonal non-coding changes near *TERT* (promoter mutations) observed in the PCAWG study (Gerstung et al., 2020). Furthermore, our results support the role of tumor initiation by *ARID1A* in many cancer types (Mathur et al., 2017; Zhai et al., 2016; Zhang et al., 2023), as well as the promotion of tumor metastasis by *ATRX* in bone cancer (Bartholf DeWitt et al., 2022).

### The effect of treatment on cancer fitness levels is stronger in primary than in metastatic tumors

Treatment is considered a strong selection pressure, substantially influencing the evolution and adaptation of various cancer types (Birkbak and McGranahan, 2020). Treatment-induced genomic alterations have been observed in many metastatic tumors, such as focal amplification of *MDM2* and copy gains of *GNRHR* have been identified in ER-positive metastatic breast cancer treated with endocrine therapy (Robinson et al., 2013). Additionally, mutations in certain genes, particularly those related to drug resistance, present treatment-induced selection pressures in cancer. Notable examples include alterations in *AR* in prostate cancer (by androgen-deprivation therapy), *ESR1* in breast cancer (by endocrine therapy), and *EGFR* in lung cancer (by *EGFR* inhibitors, commonly used in non-small cell lung cancer) are known to exhibit significant differences in genomic alteration frequencies between primary and metastatic tumors due to these therapeutic conditions (Birkbak and McGranahan, 2020; Martinez-Jimenez et al., 2023; Nguyen et al., 2022). Based on these previous findings, we next investigated whether treatment could influence the fitness of cancer genes, focusing on two-way interactions across different cancer types and states, separately.

In the MSK-MET cohort, treated samples accounted for 14.88% of primary tumors (ranging from 5.73% to 22.74% across 10 cancer types) and 39.03% of metastatic tumors (ranging from 21.12% to 58.22%; **Figure 3A**). Despite metastatic tumors consistently displaying an overall increase in clonality fraction compared to their primary counterparts (observed in 7 out of 10 cancer types), the data did not show a significant influence of tumor clonality based on treatment conditions across cancer types (**Figure S6A** and **B**). We initially compared the overall differences in two-way interaction strengths between primary and metastatic tumors, depending on treatment status. Treated tumors presented higher interaction strengths compared to their non-treated counterparts across cancer types, both in primary and metastatic tumors. Notably, six cancer types (60%) showed significantly higher interaction strength in treated samples compared to non-treated samples under at least one condition: five cancer types in the CNA-loss model (Breast, Core-GI, Gynecologic, Head-and-Neck, and Skin) and three in the CNA-gain model (Core-GI, Gynecologic, and Soft-Tissue), with two cancer types overlapping at a significance level of *P* < 0.05 by the Mann–Whitney *U*-test (**Figure 3B**). The increase in effect size was notably more evident in primary tumors compared to metastatic tumors across cancer types, both in the context of CNA-loss and gain models (with average fold-change increases ranging from 0.01 to 1.35 in primary tumors and 0.01 to 0.49 in metastatic tumors). This trend is consistently observed at the subtype level (**Figure S6C** and **D**).

**Figure 3.**
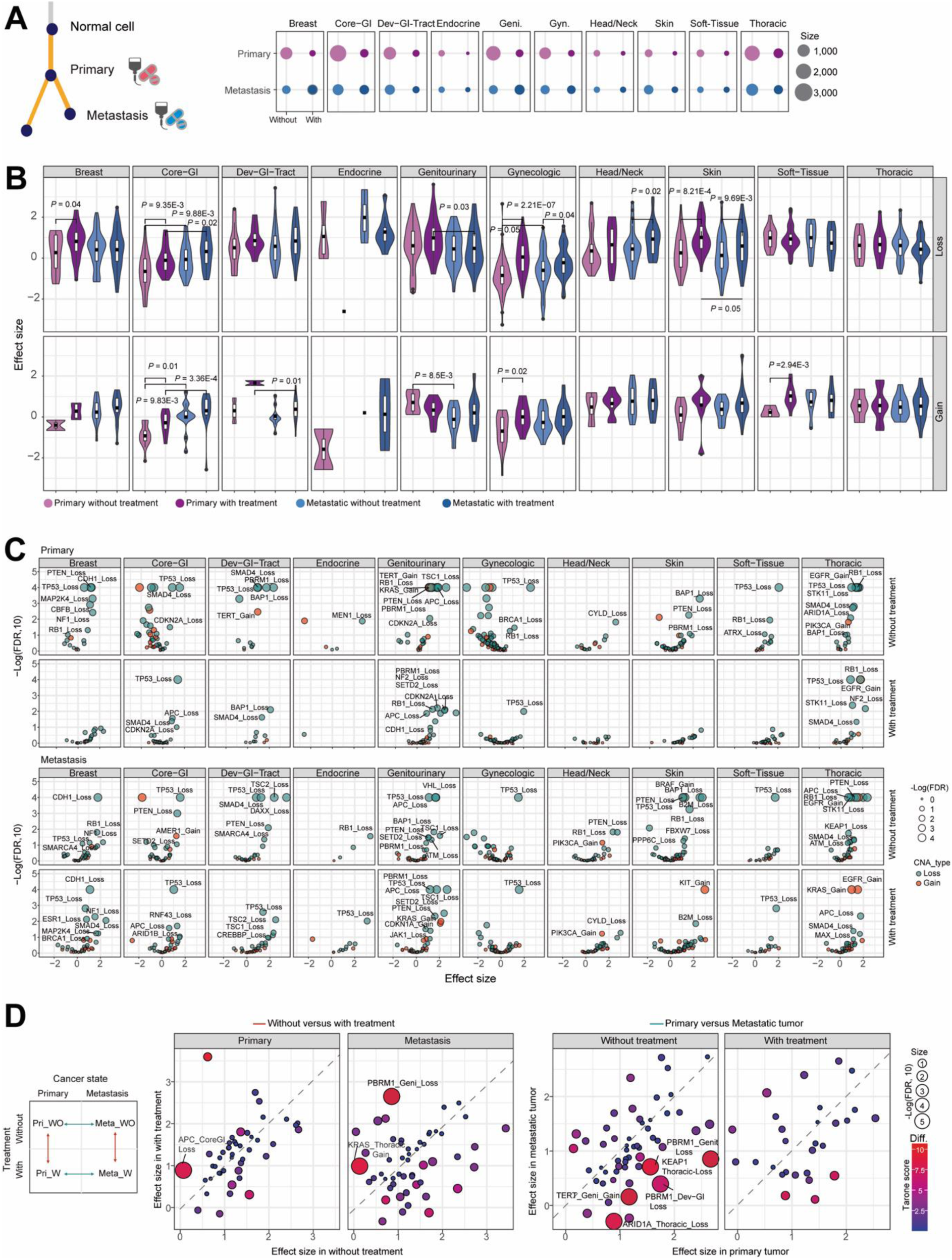
Treatment effect on two-way interaction strength across cancer states. (**a**) An overview of the size distribution of tested samples with or without treatment in each cancer type. (**b**) The average two-way interaction strength within a gene in CNA-loss (top panel) and CNA-gain (bottom panel) models across treated and non-treated conditions. Statistically significant differences across conditions were calculated by Mann–Whitney *U*-test. (**c**) Volcano plot showing the effect size of two-way interactions and their significance across various conditions. (**d**) Comparison of two-way interaction between without treatment and with treated tumors in the same cancer state (left panel; 3 significant pairs at 10% FDR) and between primary and metastatic tumors in the same treatment condition (right panel; 5 significant pairs at 10% FDR).

Next, we aimed to identify significantly interacting gene-cancer type pairs under treated and non-treated conditions across cancer types. We conducted this analysis independently for both treated and non-treated conditions (i.e., interaction in primary breast cancer under treated condition or primary breast cancer under non-treated condition), considering pairs with mutation frequencies > 1% and CNAs frequencies > 10% in each condition. Across these conditions, we tested a total of 579 pairs involving 153 genes in primary tumors (comprising 292 pairs in treated and 287 pairs in non-treated conditions) and 810 pairs involving 153 genes in metastatic tumors (consisting of 430 pairs in treated and 380 pairs in non-treated conditions). In total, we identified 70 interactions (12.09%, with 7.19% in treated and 17.07% in non-treated conditions) in primary tumors and 87 interactions (10.74%, with 8.37% in treated and 13.42% in non-treated conditions) in metastatic tumors at FDR 10%, including 35 overlapping pairs (**Figure 3C** and **Figure S7**; **Tables S6**). As expected, well-known drug-resistant genes in particular cancer types presented significant interactions between mutations and CNAs, including interactions between mutations and CNA-gain of *EGFR* in Thoracic cancer, mutations and CNA-loss of *NF1* in breast cancer, and mutations and CNA-loss of *PTEN* in Genitourinary cancer (**Figure S8A**). Next, we conducted a position-specific analysis on the identified pairs to identify the exact genomic positions that significantly contribute to these interactions (**Figure S9**), including R273C or R175H for *TP53*, and G12V or C for *KRAS*.

While we observed significantly overlapping pairs between treated and non-treated samples in both primary and metastatic tumors (odds ratio = 2.64, *P*-val = 2.67E-8 in primary tumors and odds ratio = 8.17, *P*-val = 2.86E-42 in metastatic tumors by Fisher’s one-way test), some pairs exhibited distinct interaction strength depending on the treatment condition, whether enriched in treated or non-treated samples. Notably, *APC* mutations and CNA-loss exhibited significantly increased interaction strength in primary Core-GI tumor under treated condition compared to non-treated condition (effect size in non-treated = 0.04 versus treated = 0.89; **Tables S6**). Additionally, *PBRM1* mutations and CNA-loss in metastatic genitourinary cancer exhibited markedly higher interaction strengths in treated compared to non-treated condition (effect size in non-treatment = 0.85 versus treated = 2.65; **Tables S6**). To further explore condition-specific two-way interactions, we examined whether the interaction strength of mutations and CNAs within a gene varied across two different conditions: (i) non-treated versus treated tumors within the same cancer state, and (ii) primary versus metastatic tumors within the same treatment condition, employing an odds ratio heterogeneity test (See **Methods**; **Figure 3D** and **Figure S8B** and **C**). We tested 97 pairs involving 36 genes to determine if their effect sizes differed significantly based on treatment state (e.g., treated vs. non-treated primary tumors), and 84 pairs involving 33 genes to see if their effect sizes varied significantly based on cancer state (e.g., treated primary versus treated metastatic tumors). At an FDR of 10%, three (non-treated versus treated) and five (primary versus metastatic tumors) pairs exhibited significantly different effect sizes in each comparison (**Tables S7** and **S8**), including one overlapping pair, *PBRM1* in Genitourinary tumors. At an FDR of 20%, 6 and 14 interactions were detected separately, including one additional overlapping pair, *KEAP1* in Thoracic cancer. In detail, significant interactions were found between *PBRM1* mutation and CNA-loss in Genitourinary metastatic tumors (effect size in non-treated = 0.85 versus treated = 2.65; *P* = 1.07E-3), *KRAS* mutation and CNA-gain in Thoracic metastatic tumors (0.13 versus 0.99; *P* = 4.73E-3), and APC mutation and CNA-loss in Core-GI primary tumors (0.04 versus 0.89; *P* = 6.85E-3; **Figure 3D**). Although these genes were not previously recognized as treatment resistance genes in the detected cancer types, they have been identified as potential targets, such as *KRAS* in Thoracic metastatic cancer [33]. Notably, *PBRM1* has frequent loss-of-function mutations in clear cell renal carcinoma (ccRCC) that are associated with clinical benefit from immune checkpoint therapies (Miao et al., 2018). Also, *APC*, an initiating factor for tumor progression in colon cancer and suggested as a target gene in this cancer type (Vande Voorde et al., 2023). Among five significant pairs between two different cancer states under the same treatment condition, *KEAP1*, as a known contributor for lung cancer tumorigenesis and metastasis (Lignitto et al., 2019; Romero et al., 2017), presented a significantly different effect size in primary non-treated condition compared to metastatic non-treated condition (effect size in non-treated primary tumors = 1.56 versus metastatic tumors = 0.71; odds ratio heterogeneity test *P*-value = 4.42E-3).

### High-order interactions are enriched in cancer hallmarks and are distinct between primary and metastatic cancer states

In diverse cellular contexts, the fitness level of a cancer gene can be modulated by other genes, particularly through high-order interactions involving multiple genomic alterations. For example, the two-way interaction between the mutation and CNAs of gene A can be affected by the mutation status of another gene B. This influence can result in either an increase (positive high-order interaction) or decrease (negative high-order interaction) in fitness of gene A. This forms a three-way interaction, also known as a high-order interaction (Domingo et al., 2019; Taylor and Ehrenreich, 2015). In melanoma, a notable example highlights a strong two-way interaction between *BRAF* mutations and *BRAF* copy number gain, which is observed in samples without *NRAS* mutations but decreased in samples carrying *NRAS* mutations (Park et al., 2021). It is known to play a pivotal role in shaping various phenotypic traits from model organism to human (Costanzo et al., 2019; Kuzmin et al., 2018), including drug resistance, genetic diseases, and complex diseases (Celaj et al., 2020; Cordell, 2009; Lozovsky et al., 2021; Park et al., 2021). Therefore, understanding the complexity of these interactions is crucial for enhancing cancer drug discovery and development (Ryan et al., 2023). For example, mutations in *PTEN* in breast tumor cell lines have been associated with a synergistic response to a combination of PI3K and AKT inhibitors, as revealed through extensive drug screening involving approximately 2,000 drug combinations (Jaaks et al., 2022).

To identify cancer-state-specific high-order interactions and understand the corresponding fitness landscape, we used a linear regression model to analyze interactions between three genomic alterations across cancer types, separately for each cancer state. This approach aimed to detect high-order interactions where the two-way interaction (mutation and CNAs within gene A) is perturbed based on the mutation status of gene B (**Figure 4A**; see **Methods**). Initially, we compiled a set of 124 significant two-way interactions involving 47 genes to identify high-order interactions (**Tables S9**). This set comprised 51 interactions from 27 genes observed in primary tumors and 73 interactions from 42 genes in metastatic tumors, with an overlap of 34 interactions. Systematic analysis was conducted on each pair (i.e., gene A two-way interaction) to detect higher-order interactions, incorporating mutations from an average of 47.45 other cancer genes (median = 41, with at least 1% mutation frequency in that condition; min = 11, max = 99) as a third alteration (i.e., gene B) in primary tumors, and an average of 40.97 other cancer genes (median = 39; min = 10, max = 70) in metastatic tumors. In total, three-way interactions involving two genes and their three genomic alterations were tested in 2,420 pairs for primary and 2,991 for metastatic tumors.

We identified 59 high-order interactions from 22 genes, including 38 interactions in primary tumors and 21 interactions in metastatic tumors, with 1 overlapping pair (*TP53* mutations, *TP53* CNA-loss, and *TERT* mutations in Genitourinary tumors) at FDR 10% (**Figure 4B-C** and **Figure S10**). The majority of high-order interactions (77.97%, 46 out of 59; 76.32% in primary and 80.95% in metastatic tumor) exhibited a decreased interaction strength between mutations and CNAs of the first gene in the presence of a mutation in a second gene as we previously observed with TCGA study (Park et al., 2021). Notably, many of the third-order interactions, both from primary and metastatic tumors, significantly involved cancer hallmarks, including Cell cycle/apoptosis and MAPK pathway (Li et al., 2023). This indicates that these genes play crucial roles in fundamental cancer functions and that third-order interactions share associated functional roles in cancer. This suggests a fundamental principle in cancer genomics: two genes connected by negative three-way interactions tend to be partially redundant, requiring activation either through two alterations within a single gene or between the two genes. Interestingly, as previously mentioned, there was a substantial overlap in two-way interactions between primary and metastatic tumors with 34 overlapping interactions (out of 51 in primary and 73 in metastatic tumors). However, only one three-way interaction was detected in both primary and metastatic tumors (interaction between *TP53* mutation, *TP53* CNA-loss, and *TERT* alterations), highlighting the cancer-state-specific characteristics of high-order interactions. Consistently, three-way interactions from primary and metastatic tumors exhibited functional annotations that are specifically enriched for each cancer state (Fisher’s exact test, *P* < 0.05; **Figure S11**).

### Interaction between mutations and CNAs is associated with patient survival time

To explore how different genetic interactions within a gene or between genes contribute to cancer progression and verify their clinical implications, we hypothesized that if a two-way interaction (co-occurrence of mutations and CNAs within a gene) has a greater effect on cancer fitness than a single genomic alteration (either a mutation alone or a CNA alone), then these co-occurrences within a gene would have a stronger influence on patient survival than a single alteration. To test this hypothesis, we conducted a systematic survival analysis of detected two-way interacting pairs (*N* = 124) using clinical information, comparing the prognosis of cancer patients with both mutations and CNAs to those with either a mutation only or a CNA only (**Figure 5A**).

**Figure 5.**
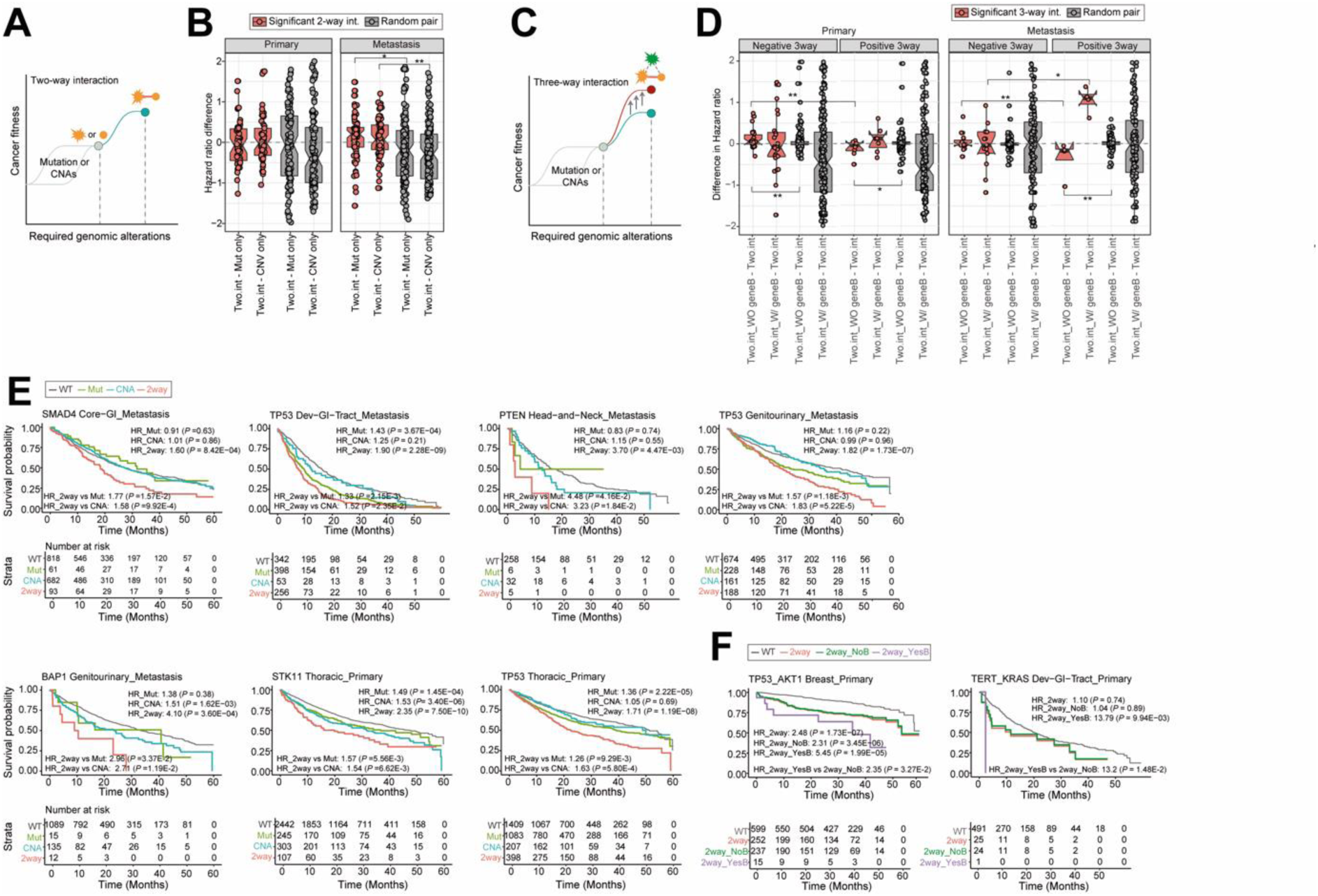
Impact of genomic alterations within and between genes on cancer patient prognosis. (**a**) Model for two-way interaction in cancer fitness level. In the two-hit driver gene model, a single genomic alteration (either a mutation or CNA) can increase fitness levels. A combination of two genomic alterations, though, is more effective and necessary to achieve optimal fitness. (**b**) Hazard ratio (HR) differences between samples with co-occurrence of mutations and CNAs within a gene and samples with single-genomic alterations were analyzed. Specifically, 124 two-way interacting pairs were compared with 526 random, non-significant pairs. The differences in hazard ratios between samples with co-occurrences of two-genomic alterations (i.e., two-way interactions) and single-genomic alterations (either mutation only or CNA only) are presented. For the box plots, the center black line represents the median. A multivariate cox regression model was applied to measure the hazard ratio. (**c**) Impact of additional genomic alterations on two-way interaction strength. While the two-way interaction between genomic alterations within a gene affects cancer fitness, the mutation status of another gene can modulate this interaction’s strength. (**d**) HR differences of samples with co-occurrence of mutation and CNAs within the first gene, compared to samples with or without mutations of a second gene. The two-way interaction strength can increase (positive interaction; *N*=13) or decrease (negative interaction; *N*=46) depending on the mutation status of the second gene. Random interactions (non-significant three-way interacting pairs; *N*=1,645) were used as controls. The center black line in the boxplots represents the median. HRs were measured using a multivariate Cox regression model. (**e**) Kaplan-Meier curves comparing the overall survival outcomes of cancer patients based on their genomic alteration status, with colors representing different patient groups. To specifically measure the contribution of 2-way interactions, HR of samples with co-occurrences of two genomic events, compared to samples with either mutation or CNA, are also presented. The plot uses a maximum five-year survival period. HRs and corresponding *P*-values from the Wald test are presented. Detailed HR, 95% confidence intervals, and *P*-values are summarized in **Tables S10**. (**f**) Kaplan-Meier curves comparing the overall survival outcomes of cancer patients based on their genomic alteration status, with colors representing different patient groups. To measure the contribution of 3-way interaction specifically, HR of samples with co-occurrences of two-genomic events with or without mutation of the second gene. Maximum five-year survival was used in this plot. HRs and corresponding *P*-values from Wald test are presented. Detailed HR, 95% confidence interval, and *P*-value are summarized in **Tables S11**.

As expected, many detected two-way interacting pairs show significantly high hazard ratios (31 pairs: 14 out of 51 in primary tumors and 17 out of 73 in metastatic tumors, with hazard ratios > 1 and Wald test *P*-value < 0.05; representing frequencies of 27.45% and 23.29%, respectively; **Tables S10**). Moreover, samples with co-occurrences of two genomic alterations had significantly higher hazard ratios than samples with only mutations (*P* = 5.38E-2 in metastatic tumors, Mann–Whitney *U*-test) and samples with only CNAs (*P* = 1.18E-3 in metastatic tumors) when compared to random pairs (i.e., without significant co-occurrence between mutation and CNAs within a gene; **Figure 5B**). This indicates that samples with co-occurrence of two genomic events have a worse prognosis (i.e., shorter overall survival time) than samples with single-genomic alterations. After applying the stringent condition that the hazard ratio for the two-way interaction is significantly higher under both the separate mutation-only and CNA-only conditions (i.e., not based on the wild-type sample, Wald test *P*-value < 0.05), seven pairs were identified (**Figure 5E**). For example, Genitourinary metastatic patients harboring both *BAP1* mutations and *BAP1* CNA-loss, detected as a two-way interacting pair (effect size = 1.77, *P*-val = 4.04E-6), exhibited a significantly stronger decrease in overall survival (average overall survival time = 9.69 months; hazard ratio = 4.10, 95% confidence interval [CI]: 1.89-8.89, Wald test *P*-val = 3.60E-4) than patients with only *BAP1* mutations (24.14 months; hazard ratio = 1.38, 95% CI: 0.67-2.84, *P*-val = 0.38) and patients with only *BAP1* CNA-loss (21.28 months; hazard ratio = 1.51, 95% CI: 1.17-1.96, *P*-val = 1.62E-3; **Figure 5E**). This worse prognosis in samples with co-occurrence of two genomic events remains robust even after accounting for the *BAP1* mutation background (hazard ratio = 2.96, 95% CI: 1.09-8.07, *P*-val = 3.37E-2) or *BAP1* CNA-loss background (hazard ratio = 2.71, 95% CI: 1.25-5.88, *P*-val = 1.19E-2).

Furthermore, we extended this hypothesis to three-way interactions. Since we proposed the evidence of three-way interactions that could modulate the strength of two-way interaction depending on the mutation status of another gene, we expected that prognosis of cancer patients could be further influenced by the presence of three-way interactions (**Figure 5C**). Consistent with our hypothesis, many of the 59 detected negative three-way interaction pairs show significantly higher hazard ratios in samples with two-way interactions in the first gene but without mutations in the second gene (19 pairs: 15 out of 29 in primary tumors and 4 out of 17 in metastatic tumors, with hazard ratios > 1 and Wald test *P*-value < 0.05; frequencies of 51.72% and 23.53%, respectively). Conversely, many positive three-way interacting pairs show significantly high hazard ratios in samples with two-way interactions in the first gene and mutations in the second gene (6 pairs: 3 out of 9 in primary tumors and 3 out of 4 in metastatic tumors, with hazard ratios > 1 and Wald test *P*-value < 0.05; frequencies of 33.33% and 75.0%, respectively). The full list of pairs is provided in **Tables S11**. This indicates that negative three-way interactions tend to have higher hazard ratios in samples with two-way interactions without the mutation of another gene compared to positive three-way interactions (*P* = 2.90E-3 in primary and *P* = 3.01E-3 in metastatic tumor, Mann–Whitney *U*-test) and random interaction pairs (*P* = 8.86E-4 in primary; **Figure 5D**). Conversely, positive three-way interactions tend to show higher hazard ratios in samples with two-way interactions that include mutations in another gene compared to negative interactions (*P* = 1.83E-2 in metastatic tumors). After applying the stringent condition that the hazard ratio is significantly higher when comparing the two-way interaction with and without the second gene mutation (Wald test *P*-value < 0.05), two pairs were selected (**Figure 5F**). In particular, there is a strong third-order interaction between *TP53* mutations, *TP53* CNA loss, and *AKT1* mutations in primary breast cancer (effect size of three-way interaction = 1.65, *P*-val = 5.06E-3). Although samples with the two-way interaction between *TP53* mutation and *TP53* loss show a worse prognosis (average overall survival time = 28.62 months versus 35.52 months in wild-type samples; hazard ratio = 2.48, 95% CI: 1.76-3.48, Wald test *P*-val = 1.73E-7), samples with an additional *AKT1* mutation exhibit much shorter survival times (average overall survival time = 25.35 months; hazard ratio = 5.45, 95% CI: 2.50-11.87, Wald test *P*-val = 1.99E−05; **Figure 5F**). This worse prognosis in samples with the *AKT1* mutation remains robust even after accounting for the *TP53* two-way interaction without the *AKT1* mutation background (hazard ratio = 2.35, 95% CI: 1.07-5.17, *P*-val = 3.37E-2). Collectively, these analyses demonstrate how genetic and high-order interactions contribute to tumorigenesis, and how the optimal combination of genetic alterations within or between genes enhances cancer fitness, impacting the prognosis of cancer patients.

## Discussion

Our study is the first to undertake a large-scale analysis of approximately 25,000 samples, investigating whether the optimal cancer fitness level (i.e., number of genomic events a cancer gene requires) can be changed based on the state of the cancer. Notably, a portion of cancer genes robustly exhibit cancer-state-specific fitness level changes (referred to as the perturbed group), which may play critical roles in the transition from primary to metastatic tumors. We have also demonstrated that optimal cancer fitness levels can change depending on treatment status across cancer states and types, suggesting these could be cancer-state-specific target genes for therapies. Additionally, we have identified cancer-state-specific three-way interactions across cancer types, which not only play essential roles in tumorigenesis but also reflect cancer-state-specific functions. Indeed, we have provided multiple examples where two-way or three-way interactions impact overall survival times to patients, underscoring the clinical significance of these interactions.

The primary objective of this study was to explore the interplay between mutation and copy-number alterations within a cancer gene, elucidating their potential differential impacts across two different cancer states. While this two-way interaction constitutes the major focus, it’s imperative to acknowledge additional factors such as epigenetic modifications (e.g., promoter DNA hypermethylation) or alterations in gene expression (Terekhanova et al., 2023). Although epigenomic and transcriptomic data were not available in the MSK-MET dataset to our knowledge, several studies have shed light on cancer-state specific differences beyond the genomic alterations (Garcia-Recio et al., 2023; Nishiyama and Nakanishi, 2021). For instance, a study involving 55 females with metastatic breast cancer revealed that 33.3% of cases exhibited gene expression-based subtypes switching distinctly between primary and metastatic tumors, underscoring cancer-state specific changes in gene expression (Garcia-Recio et al., 2023). Furthermore, HLA-A promoter methylation exhibited a significantly higher prevalence in metastatic tumors compared to primary tumors, potentially influencing the tumor microenvironment by attenuating immune cell infiltrations (Garcia-Recio et al., 2023). These findings emphasize the necessity of considering non-mutational mechanisms and the tumor microenvironment for a comprehensive understanding for the differences between primary and metastatic tumors.

While MSK-MET stands as one of the largest datasets of cancer genomes to date, featuring a balanced representation of primary and metastatic patients via a consistent pipeline, several limitations still need to be explored in this study. Our number of significant interactions in two-way model in primary or metastatic tumors and dividing them by treatment conditions is still not saturated and is expected to be increased in power as sample sizes increase from the saturation analysis. This indicates that possible un-identified interactions remain, and more samples will be still required to gather and understand the complete view (**Figure S4A** and **6E**). Next, MSK-MET data was generated from gene panel sequencing, which covers a defined set of cancer genes and not the whole exome or genome. As a result, the full range of genomic interactions based on cancer states may not be accurately captured. For example, in our analysis, clonal fractions between treated and non-treated samples were not significantly different (**Figure S6A** and **B**). Conversely, several other whole-exome or genome sequencing-based studies have observed significant differences (Hu et al., 2020; Martinez-Jimenez et al., 2023), suggesting the potential limitations of a gene-panel-based approach. Additionally, our primary and metastatic samples were not paired (i.e., they originated from different patients) which will be crucially necessary for the cancer evolutionary analysis. Resources such as the TRAcking Cancer Evolution through therapy (TRACERx) consortium (Al Bakir et al., 2023; Mitchell et al., 2018; Turajlic and Swanton, 2016; Turajlic et al., 2018a; Turajlic et al., 2018b) — valuable complementary resource to matched patients after tracking many years with therapeutic implications covering lung, skin, kidney, and prostate — could potentially offer complementary insights. Last, details regarding treatment information (such as types of treatment, duration of treatment, etc.) were not provided in the original resource, thus limiting the understanding of treatment type-specific cancer fitness changes.

In conclusion, our findings underscore the role of differential cancer fitness levels in contributing to cancer states between primary and metastatic tumors. By measuring the associations between mutation and copy-number changes within a gene, we provide a potential new perspective to understand cancer state differences that extends beyond current approaches focusing on differences in single-genomic differences. Our study suggests that relying solely on single-genomic alterations may not suffice to comprehend the distinctions between primary and metastatic cancer states. As such, our approach forms a valuable methodology that could potentially be adapted to further study other aspects of tumor progression.

## Acknowledgements

The results shown here are in whole or part based upon data generated by the MSK-MET. We thank Luis Garcia-Jimeno for assistance with log-regression model. S.P. is supported by the Agencia Estatal de Investigación, Ministerio de Ciencia e Innovación (MCIN/AEI/10.13039/501100011033) through the RETOS project PID2019-109571RA-I00. S.P. is also supported by Ramón y Cajal, Ramón Areces Foundation and Agencia Estatal de Investigación. S.P. acknowledges the support of the Severo Ochoa Centres of Excellence program to the CNIO (MCIN/AEI/10.13039/50110001103).

## Author contributions

Conceptualization— LOM, AMR, and SP; data curation— LOM, AMR, and SP; formal analysis— LOM, AMR, and SP; funding acquisition—SP; investigation—SP; methodology— LOM, AMR, and SP; project administration—SP. Resources— LOM, AMR, and SP; supervision—SP; validation— LOM, AMR, and SP; roles/writing–original draft — LOM, AMR, and SP; Writing–review and editing—all.

## Supplemental information

**Document S1. Figures S1–S12**

**Table S1. Excel file containing additional data too large to fit in a PDF, related to Table 1**

**Table S1. List of 10 cancer types and 50 sub-types in the MSK-MET cohort.**

**Table S2. Analysis of two-way interactions within cancer genes across cancer types and cancer states.**

**Table S3. Cancer gene classification based on two-way interaction preference between primary and metastatic tumors.**

**Table S4. Analysis of two-way interactions within cancer genes after dividing primary samples into primary without metastasis and with metastasis.**

**Table S5. Cancer gene classification based on two-way interaction preference across different cancer states (primary without metastatsis, primary with metastasis).**

**Table S6. Analysis of two-way interactions within cancer genes across different cancer types and states, with samples categorized into treatment groups.**

**Table S7. Odds ratio heterogeneity analysis comparing treated and non-treated samples across different cancer types and states.**

**Table S8. Odds ratio heterogeneity analysis comparing primary and metastatic samples across different cancer types and treatment conditions.**

**Table S9. Analysis of three-way interactions within cancer genes across cancer types and cancer states.**

**Table S10. Survival analysis with two-way interacting pairs.**

**Table S11. Survival analysis with three-way interaction pairs.**

## STAR Methods

### Cancer genome data

In this study, we measured the interactions between two genomic alterations (mutations and copy-number alterations) within cancer genes. Our analysis was conducted using the MSK-MET dataset (Nguyen et al., 2022). To ensure the use of a high-quality dataset, rigorous exclusion criteria was already applied to the original dataset, eliminating samples that exhibited low sequencing coverage, low tumor purity, pediatric patients, cancers with unknown primary origins, or those lacking available matched normal samples. As a result, our dataset comprises comprehensive molecular data and clinical outcome information pertaining to a substantial cohort of 25,775 patients. These patients provided unmatched samples, including primary tumors (15,632 samples, 60.65%) and metastatic tumors (10,143 samples, 39.35%), encompassing 10 cancer types. The clinical data was directly sourced from the MSK-MET dataset, encompassing key patient attributes such as age at the time of sequencing (available for 99.0% of samples), sample type (distinguishing between primary and metastasis), cancer type and sub-type (curated subtype information), treatment conditions (prior treatment status was available for 100% of the samples), the number of metastatic events (MET_COUNT, available for 98.69% of samples), and sample quality (including sequence coverage, available for all patients, and sample purity, available for 98.42% of patients). Furthermore, we categorized primary samples into two distinct groups: primary tumors from non-metastatic patients (24.89% of primary samples; labeled as P no met with MET_COUNT = 0) and primary tumors from metastatic patients (75.11% of primary samples; labeled as P met with MET_COUNT > 0). This comprehensive dataset serves as the foundation for our in-depth exploration of cancer genomic interactions and their implications.

### Somatic alterations

We obtained genomic data from the MSK-MET dataset via cBioPortal (https://www.cbiopor-tal.org/study/summary?id=msk_met_2021;data_mutations.txt). This dataset encompasses 230,419 functional variants, including premature truncation mutations, non-synonymous mutations, and single-residue substitutions for 492 genes. To determine the clinical actionability of somatic mutations, we utilized the Oncology Knowledge Base (OncoKB) (Chakravarty et al., 2017) in conjunction with MafAnnotator v.3.3. Only variants classified as “Oncogenic” or “Likely Oncogenic” were considered, resulting in the inclusion of 37.0% of the initial somatic functional variants (85,262 out of 230,419 variants). Additionally, we incorporated upstream gene variants related to *TERT* based on OncoKB annotations. To assess the clonality of each variant (either clonal or sub-clonal), we referred to the facets-suite (https://github.com/mskcc/facets-suite). Of the total detected variants, 98.38% (226,680 out of 230,419) variants were categorized as either clonal (56.56%, 128,215 variants) or sub-clonal (19.87%, 45,044 variants), with the remaining variants classified as indeterminate (8.72%, 19,766 variants) or not available (14.85%, 33,655 variants). Furthermore, we conducted a comparative analysis of tumor clonality with respect to metastatic biopsy locations across different cancer types, revealing no discernible differences (**Figure S12**). Arm-level copy-number alterations (either loss or gain) were downloaded from cBioPortal (data_cna_hg19.seg), calculated using the ASCETS tool (Nguyen et al., 2022).

### Somatic driver genes

To collect high-confidence cancer genes, we relied on MSK’s precision OncoKB database (retrieved in September 2022) (Chakravarty et al., 2017). A total of 568 genes were provided, categorized as Tumor Suppressor Genes (TSGs, *N*=268), Oncogenes (OGs, *N*=261), and Dual-Functional Genes (DFGs, *N*=39) based on mutation type (activation or inactivation) and frequencies of loss-of-function/copy-number deletions (Davoli et al., 2013; Vogelstein et al., 2013). DFGs were defined when genes could be classified as both TSGs and OGs. In total, 384 genes (195 TSGs, 167 OGs, and 22 DFGs) exhibit somatic mutations (likely/oncogenic variants) or copy-number alterations (CNAs) in the MSK-MET dataset, thus qualifying for interaction testing across ten different tissues. Additionally, we selected genes in cancer types with mutation frequencies > 1% and CNAs > 10% (separately for Loss and Gain models) for testing co-occurrence between mutations and CNAs within a gene. This process resulted in the assignment of 299 pairs for 126 genes in the primary state (232 pairs for 101 genes in the Loss model, 67 pairs for 36 genes in the Gain model, with 5 pairs for 5 genes tested in both models). In the metastatic state, 351 pairs were assigned for 120 genes (241 pairs for 99 genes in the Loss model, 110 pairs for 60 genes in the Gain model, with 30 pairs for 25 genes tested in both models).

### Binary matrix preparation

The final set of binary calls for genomic alterations, encompassing somatic mutation calls and CNAs (loss/deletion or gain/amplification), was assigned to genes across samples. Somatic mutation calls were assigned ’1’ when the gene harbored at least one clinically actionable somatic mutation, while CNAs were assigned ’1’ when the gene exhibited loss (for CNA-Loss) or gain (for CNA-Gain). After defining these binary calls for the 384 genes, we excluded 355 samples from the final analysis as they showed no detectable genomic alterations, leaving us with a dataset of 25,420 samples across ten cancer types (https://github.com/SolipParkLab/PriMet_Fitness).

### Statistical model for mutation-CNAs associations

We assessed the two-way interactions between somatic mutations and CNAs (either CNA-Loss or CNA-Gain) within genes across cancer types with a log-linear regression model, utilizing R (v.4.2.2), as detailed in our previous study (Park et al., 2021). This analysis involved two separate models for each gene-tissue pair:

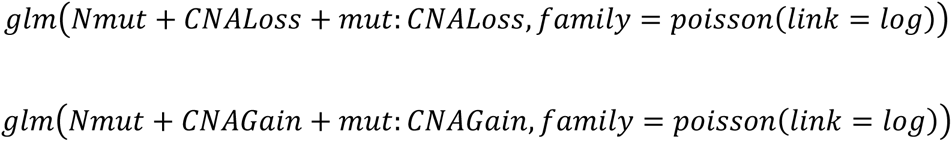

In these models, the variables are defined as follows: *mut* represents the number of samples with somatic mutations in the tested gene, *CNA Loss* (or *CNA Gain*) represents the number of samples with CNA Loss (or Gain) in the tested gene, and *mut:CNALoss* (or *mut:CNAGain*) represents the number of samples with both a mutation and CNA Loss (or Gain) in the tested gene. The strength of the two-way interaction between somatic mutation and CNAs was determined by comparing samples with CNA wild-type (no copy-number changes) to samples with CNA Loss (or Gain). To ensure robust calculations, a pseudo count of 1 was added to each frequency.

We computed regression coefficients (interaction effect sizes) and *P*-values for each gene-tissue pair individually based on *mut:CNA* interactions, utilizing the summary function in R. Positive values indicate stronger co-occurrences between mutation and CNA within the tested pair. To assess the significance of two-way interactions, we applied a permutation strategy, to control for genomic alteration heterogeneity. This approach has already been proven to successfully identify the association between mutations and CNAs within the same gene using TCGA Pan-Cancer data (Park et al., 2021), as well as epistatic interactions between cancer genes (Park and Lehner, 2015). Permutated genomic alteration matrices were generated using the *permatswap* function in the R package vegan (2.6.4) (http://vegan.r-forge.r-project.org). These matrices maintained the total number of genomic alterations (somatic mutation, CNA Gain, and CNA Loss, separately) across samples, as well as the total number of alterations per sample. For specific conditions, such as statement-specific or treatment-specific models, we applied the permutation to each condition separately. After 100 permutations, we estimated the false-discovery rate (FDR) as the ratio between the number of detected interactions in each permutated matrix (i.e., false interactions) and the number of detected interactions in the observed matrix (i.e., true interactions) at each *P*-value cut-off. Additionally, to ensure the robustness of our two-way interactions, we considered possible confounding factors, including sequence coverage, sample purity, patient age, and sample clonality. However, our analysis revealed no clear evidence of confounding effects (**Figure S2 A-D**).

### Statistical model for three-way interactions between two genes

The three-way model quantifies the strength of interactions of two genomic alterations within a gene depending on the background alteration (mutation) of another gene in a specific cancer type (Park et al., 2021). This analysis involves two separate models:

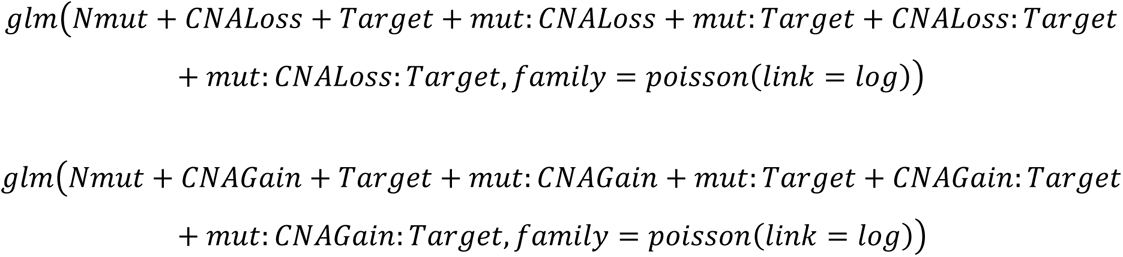

In these models, the variables are defined as follows: *mut* represents the number of samples with somatic mutations of *gene A*, *CNA* represents the number of samples with CNA of *gene A*, *Target* represents the number of samples with somatic mutations of *gene B*, *mut:CNA* represents the number of samples with both a mutation and CNA of *gene A*, *mut:Target* represents the number of samples with both a mutation of *gene A* and *gene B*, *CNA:Target* represents the number of samples with CNA of *gene A* and somatic mutation of *gene B*, and *mut:CNA:Target* represents the number of samples with somatic mutation of *gene B* when samples exhibit both a mutation and CNA of *gene A*. Regression coefficients and *P*-values are computed for pairs of two genes within a cancer type individually, derived from *mut:CNA:Target* using the summary function in R. With 100 permutations, we estimate FDR for the three-way interactions (*mut:CNA:Target*) as the ratio between the number of detected interactions in the permuted matrix (i.e., false interactions) and the number of detected interactions in the real data (i.e., true interactions) for each *P*-value cut-off (**Figure S10**).

### Saturation analysis

To determine whether our analysis has reached a saturation point regarding the number of significant interactions across conditions, we conducted a subsampling analysis (**Figure 4A** and **6E**). We progressively increased the number of samples (*N*=200) in each interval until reaching the final number of samples across conditions, including primary and metastatic samples, as well as primary-non-treated, primary-treated, metastatic-non-treated, and metastatic-treated samples. Furthermore, robustness of gene set classification were also confirmed by saturation analysis (**Figure 4B**). This process was repeated 100 times for robustness.

### Odds ratio heterogeneity test

To evaluate whether two-way interactions vary under different conditions, we focused on two groups:

i. quantifying the difference between untreated and treated tumors within the same cancer state, and
ii. quantifying the difference between primary and metastatic tumors within the same treatment condition. Pairs were considered if they were significantly detected in at least one condition (e.g., either treated or non-treated, at FDR 10%) and testable in both conditions (i.e., mutation frequency > 1% and CNV frequency > 10 in both conditions). For the first group, we tested the treatment-specific condition on 97 pairs. This included 45 pairs in primary tumors (19 pairs significant in primary with treatment, 43 pairs significant in primary without treatment, and 17 overlapping pairs) and 52 pairs in metastatic tumors (27 pairs significant in metastatic with treatment, 44 pairs significant in metastatic without treatment, and 19 overlapping pairs). For the second group, we tested the state-specific condition on 84 pairs. This included 28 pairs in treated tumors (16 pairs significant in primary treated, 21 pairs significant in metastatic treated, and 9 overlapping pairs) and 56 pairs in non-treated tumors (46 pairs significant in primary non-treated, 36 pairs significant in metastatic non-treated, and 26 overlapping pairs). In each pair, we quantified the strength of two-way interactions between mutations and CNAs within a gene using a 2 × 2 contingency table, as follows:

| *With treatment* | Mutation Present | Mutation Absent |
| --- | --- | --- |
| CNA present | *CO treated* | *Aonly treated* |
| CNA absent | *Bonly treated* | *Neither treated* |

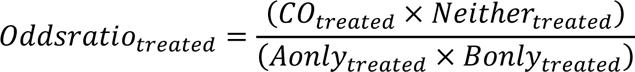

$$Oddsratio_{treated} = \frac{(CO_{treated} \times Neither_{treated})}{(Aonly_{treated} \times Bonly_{treated})}$$
| *Without treatment* | Mutation Present | Mutation Absent |
| --- | --- | --- |
| CNA present | *CO non-treated* | *Aonly non-treated* |
| CNA absent | *Bonly non-treated* | *Neither non-treated* |

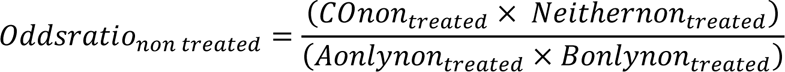

To assess odds ratio heterogeneity between the two conditions (non-treated versus treated and primary versus metastatic tumors) for each pair, we utilized Tarone’s test for quantifying the odds ratio heterogeneity. We employed the R package metaphor (version 4.4.4; https://cran.r-project.org/web/pack-ages/metafor/) for this analysis. To account for mutational heterogeneity, we compared the observed Tarone’s heterogeneity test statistics with a distribution of random statistics generated through permutation, as described above. The empirical *P*-value was calculated as the proportion of events where the random statistic exhibited a higher value than the observed statistic. We further applied the Benjamini and Hochberg method for multiple hypothesis correction to ensure the validity of our findings.

### Essential genes

We compiled a comprehensive list of 2,228 essential genes by consolidating data from two distinct sources. Firstly, we included 2,134 essential genes identified through rigorous CRISPR-Cas9 screening, characterized by robust dependencies in over 90% of pan-cancer cell lines from the DepMap database (https://depmap.org/portal/download/). Secondly, we incorporated an additional set of 297 essential genes from a study conducted by Marcotte *et al*. (Marcotte et al., 2012), encompassing three cancer types and involving 72 cell lines.

### Enrichment analysis of functional annotation

To assess the functional differences between (i) consistent and perturbed group, and (ii) high-order interaction from primary and metastatic tumors, we used gProfiler (https://biit.cs.ut.ee/gpro-filer/gost)(Raudvere et al., 2019) to measure over-representation of genes in each group against the Gene Ontology (GO; http://www.geneontology.org): Biological Process (BP), Molecular Function (MF), Cellular Component (CC) database.

After removing the redundant GO terms (frequencies of overlapping between two GO terms > 0.5 and number of annotated genes in that GO term is bigger than 500), fisher’s exact one-side test has been performed for each GO term. Additionally, we accessed a set of 21 cancer hallmarks based on multiomics data, which encompassed data from over 1,000 tumor samples and was sourced from the Clinical Proteomics Tumor Analysis Consortium (CPTAC) (Li et al., 2023).

### Statistical analysis of two-way and three-way interactions on patient survival

To assess the contribution of interaction terms (two-way and three-way) in a statistical model of patient overall survival time, Cox regression and Kaplan-Meier analyses were performed for each significant pair using the R packages survival (v.3.6.4; https://cran.r-project.org/web/packages/survival/index.html) and survminer (v.0.4.9; https://cran.r-project.org/web/packages/survminer/index.html). The Cox regression model included gender, age, and cancer subtypes as covariates to adjust the regression model. Hazard ratios and 95% confidence intervals were calculated using this model. Survival analysis was represented as a Kaplan-Meier curve with statistical significance calculated using the log-rank test.

### Data availability

This study re-analysed a MSK-MET dataset (*DOI: <u>10.1016/j.cell.2022.01.003</u>*). Somatic mutation data and copy-number alteration data were downloaded from https://www.cbioportal.org/study/sum-mary?id=msk_met_2021 alongside with clinical data containing metastatic events. Treatment data was collected from Supplementary table 1B of the MSK-MET publication (*DOI: <u>10.1016/j.cell.2022.01.003</u>*). Clonality data can be obtained by contacting the corresponding author of the original publication (*DOI: <u>10.1016/j.cell.2022.01.003</u>*). A list of cancer genes was downloaded from https://www.oncokb.org/can-cerGenes. Essential genes were collected from DepMap database https://depmap.org/portal/download/. Canonical cancer signaling pathway information were obtained from the TCGA Pan-Cancer study (DOI: *<u>10.1016/j.cell.2018.03.035</u>*) and multi-omics-based cancer hallmarks were collected from the CPTAC study (DOI: <u>0.1016/j.cell.2023.07.014</u>).

### Code availability

https://github.com/SolipParkLab/PriMet_Fitness

### Resource availability

Lead contact

Further information and requests for resources or data should be directed to and will be fulfilled by Solip Park.

### Software for statistical analysis

Regression models and statistical analyses were performed in the R statistical programming environment (v.4.2.2) using stringr (v.1.5.0), tidyr (v.1.3.1), dplyr (v.1.1.4), purr (v.1.0.2), reshape2 (v.1.4.4), tibble (v.3.2.1), survival (v.3.6.4), survminer (v.0.4.9), metafor (v.4.4.0), stats (v.4.2.2) and vegan (v.2.6.4) libraries. Figures were generated using ggplot2 (v.3.5.0), ggnewscale (v.0.4.10), ggalluvial (v.0.12.5), ggpubrr (v.0.6.0), ggrepel (v.0.9.5), ggsignif (v.0.6.4), gridExtra (v.2.3) and pheatmap (v.1.0.12). MAF file was annotated using Python (v.3.11.0) according to the premises in https://github.com/oncokb/oncokb-annotator. Network figures were generated using Cytoscape (v.3.8.0).

