## Supplementary Figures for "Mapping cancer gene dynamics through state-specific interactions"

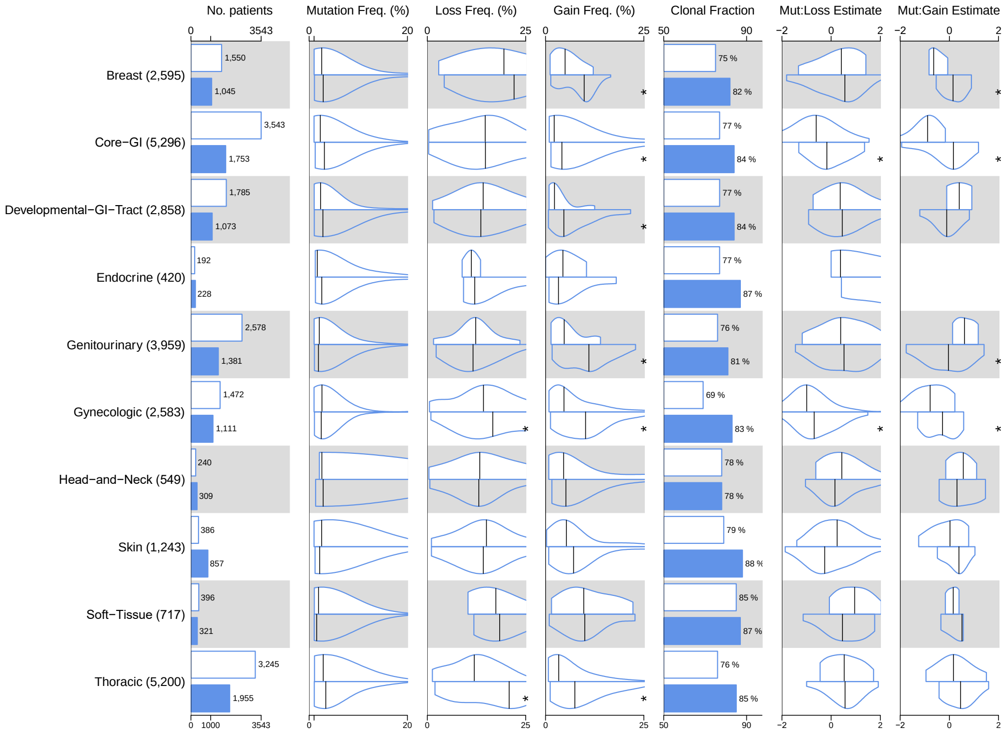


**Figure S1**. Genomic differences between primary and metastatic tumors across 10 cancer types. Presented from left to right are the following features: number of patients, oncogenic mutation frequency (%), CNA-loss frequency (%), CNA-gain frequency (%), clonal fraction (%; clonal versus subclonal), Mut-CNA loss interaction strength (effect size), and Mut-CNA gain interaction strength (effect size). Values are displayed for both primary tumors (depicted in the top white box) and metastatic tumors (depicted in the bottom blue box). Genes with a mutation frequency higher than 1% and a CNA frequency higher than 10% were included in this analysis (Mann–Whitney *U*-test *P*-value < 0.05).

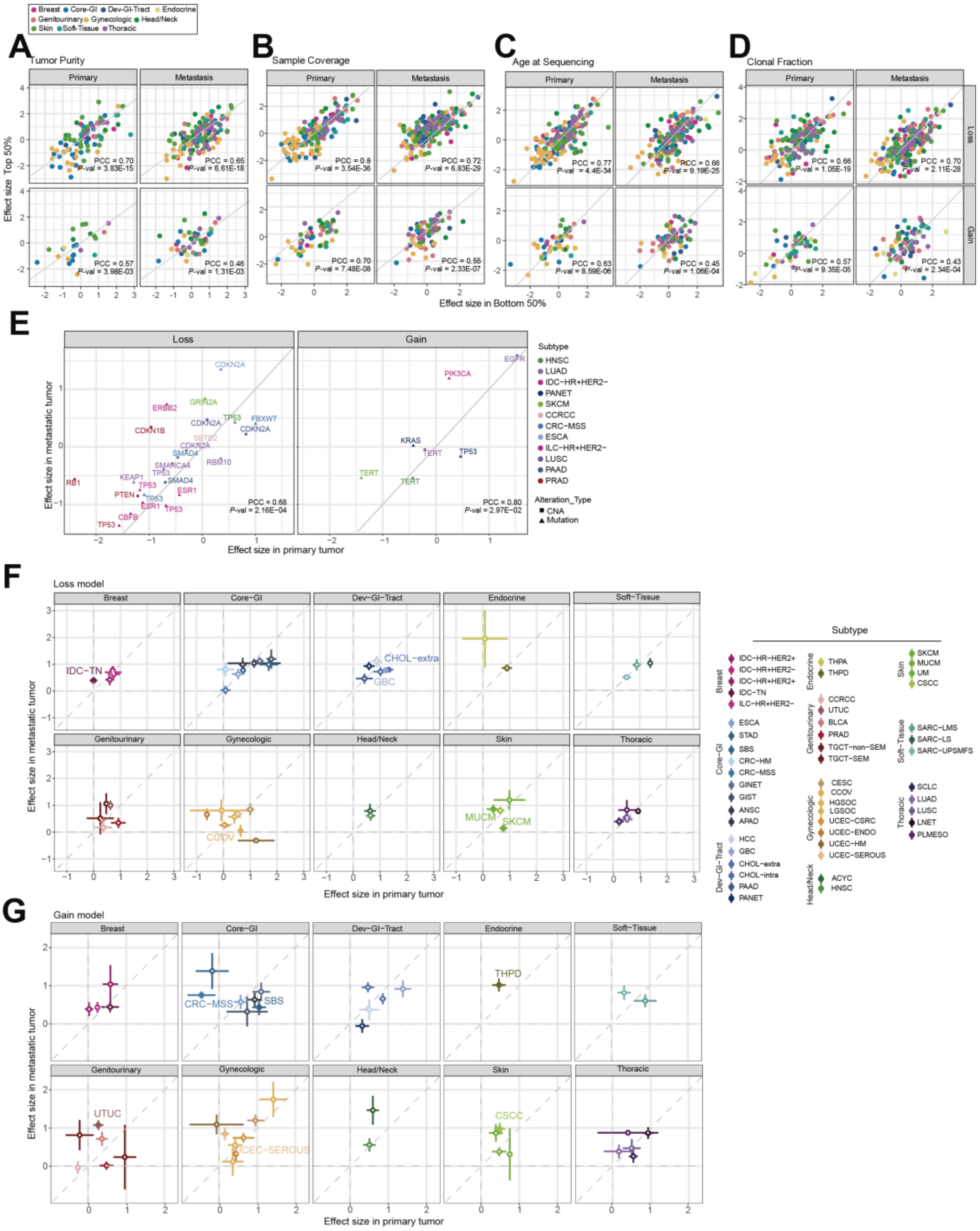


**Figure S2**. Relationship between the two-way interaction strength and various clinical/technical factors in tumor samples. The Pearson correlation coefficient (PCC) and its corresponding *P*-values are displayed. (**A**) Comparison of interaction strengths between high and low sample purity groups (samples with purity above or below the median within their respective sample states, either primary or metastasis). (**B**) Comparison of interaction strengths between high and low sample coverage groups (samples with coverage above or below the median within their respective sample states, either primary or metastasis). (**C**) Comparison of interaction strengths between high and low patient age groups (samples with patient age above or below the median within their respective sample states, either primary or metastatic). (**D**) Comparison of interaction strengths between high and low clonality groups (samples with clonality above or below the median within their respective sample states, either primary or metastatic). (**E**) Comparison of interaction strengths between primary and metastatic tumors using genes significantly differently altered across cancer subtypes (either through mutation or copy-number changes, 19 genes for 32 pairs: *Q*-value < 5% from MSK-MET original analysis (Nguyen et al., 2022)). PCCs and its corresponding *P*-values are displayed. (**F** and **G**) Effect size comparision in the subtypes level. The comparison is presented separately for two models: CNA-Loss model and CNA-Gain model. Subtypes with statistically significant differences are labelled (Mann–Whitney *U*-test *P*-value < 0.05). Error bars represent the standard error of the average effect size. For reference, abbreviations for the subtypes can be found in **Tables S1**.

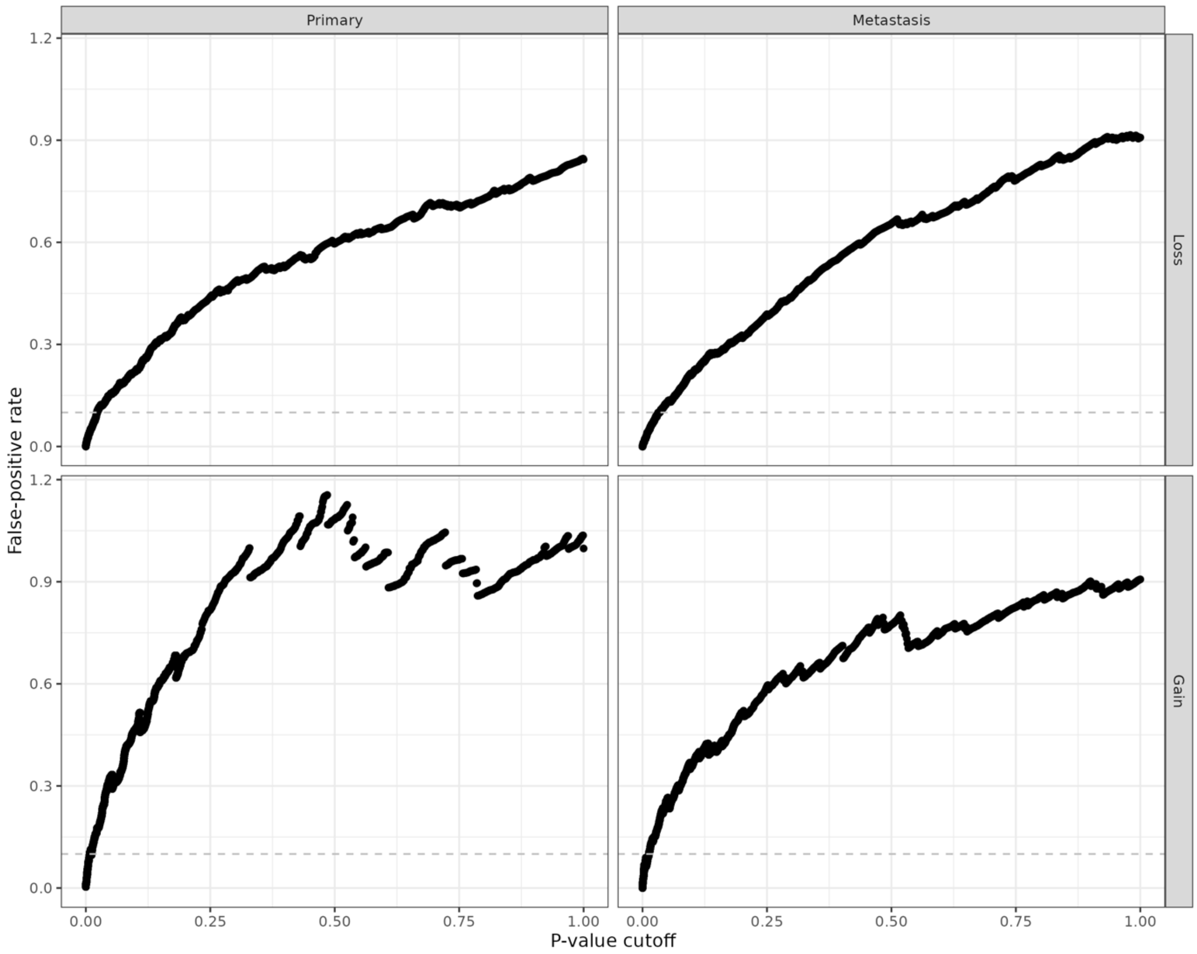


**Figure S3**. The false-positive discovery rate (FDR) values were calculated at various *P*-value thresholds using a permutation approach involving 100 iterations. This permutation strategy was employed to assess the significance of co-occurrence between pairs of genomic alterations (mutation and CNAs) within the same gene, while controlling for genomic heterogeneity within and across samples and genes. The strategy maintained the total number of alterations for each genomic event across samples as well as the total number of alterations per sample. The permutation process was performed independently for each type of genomic event (mutation, CNA-loss, and CNA-gain) and was executed separately for each cancer type. For each specified *P*-value cut-off, the FDR was estimated as the ratio between the number of detected interactions in the permuted data matrix (i.e., the false-positive interactions) and the number of interactions detected in the original data matrix (i.e., the true observed interactions). The resulting average FDR values are plotted for each *P*-value cut-off for CNA-loss (top) and CNA-gain model (bottom).

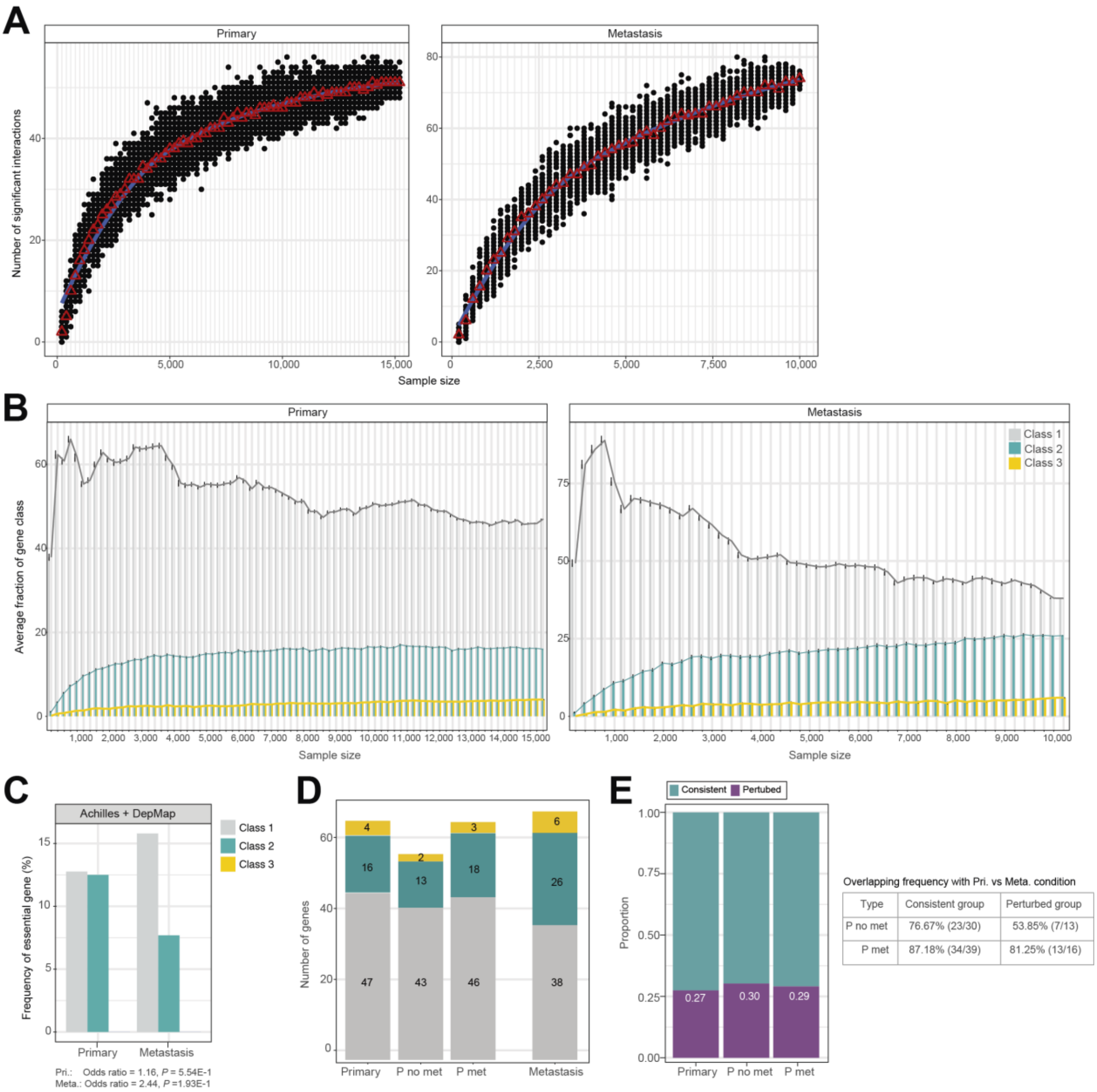


**Figure S4**. Subsampling analysis for two-way interactions-based gene classification. (**A**) Saturation analysis by subsampling tumor samples separately for primary and metastatic tumors. Each point on the graph represents a randomly selected sub-sample, with the red triangle indicating the median number of significant two-way interactions (either mut:CNA loss or mut:CNA gain) at an false discovery rate (FDR) 10%. The x-axis represents the number of samples added (every 200 samples) across 100 runs. Smoothing lines (blue) were added using the 'loess' smoothing method to visualize the number of significant interactions, and the red triangle represents the median number of significant interactions. (**B**) Average fraction of different classes of cancer genes in the detected gene-tissue pairs at varying sample sizes at an FDR 10%. Each point represents the average fraction of different classes of cancer genes, derived from randomly selected subsamples, with 200 samples added across 100 runs. Error bars indicate the standard error. **(C)** Frequency of essential genes across different gene classes in primary and metastatic tumors. Essential genes were identified through genome-wide CRISPR-Cas9 screening from DepMap (https://depmap.org/portal/download/) and Marcotte *et al* (Marcotte et al.,

2012). The odds ratio for the enrichment of Class 1 genes compared to other classes (Class 2 or 3) in essential genes, along with a one-sided Fisher’s exact test, was computed. **(D)** Different types of primary tumors to the gene classification. The distribution of genes among different gene classes when comparing various types of primary tumors and metastatic tumors. Primary tumors were categorized into those from patients without metastases (P no met) and those from patients with metastases (P met). (**E**) The proportion of consistent and perturbed groups between different types of primary tumors and metastatic tumors. Overlapping frequencies of consistent and perturbed groups were calculated between the initial comparison (i.e., primary and metastatic tumors) and different primary tumor groups (either P no met or P met).

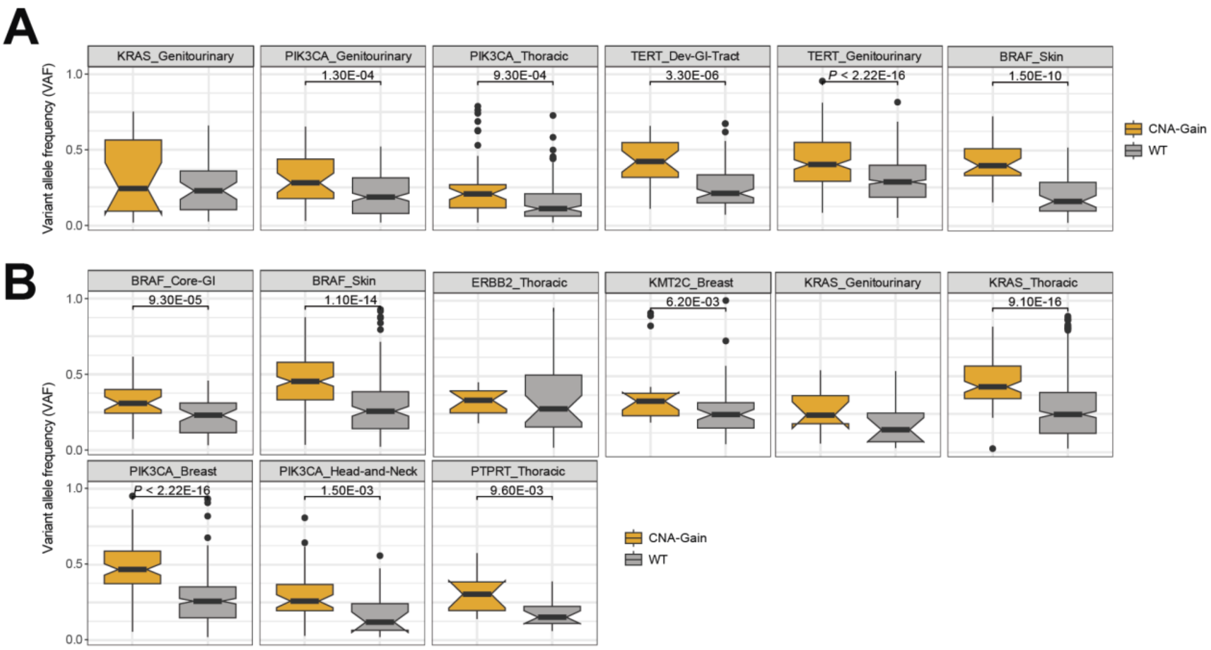


**Figure S5**. Selection for mutant allele in interactions between mutation and CNA-gain. The comparison of variant allele frequencies in significant two-way interaction pairs using (**A**) 6 pairs in primary and (**B**) 9 pairs in metastatic tumor at FDR 10%, respectively. These pairs involve mutated samples with CNA gain and CNA wild type (no CNA changes). *P*-values between two sample groups from Mann-Whitney *U* test. Each box plot shows the median value for each gene set, while the whiskers' length is 1.5 times the interquartile range, represented by the height of each box.

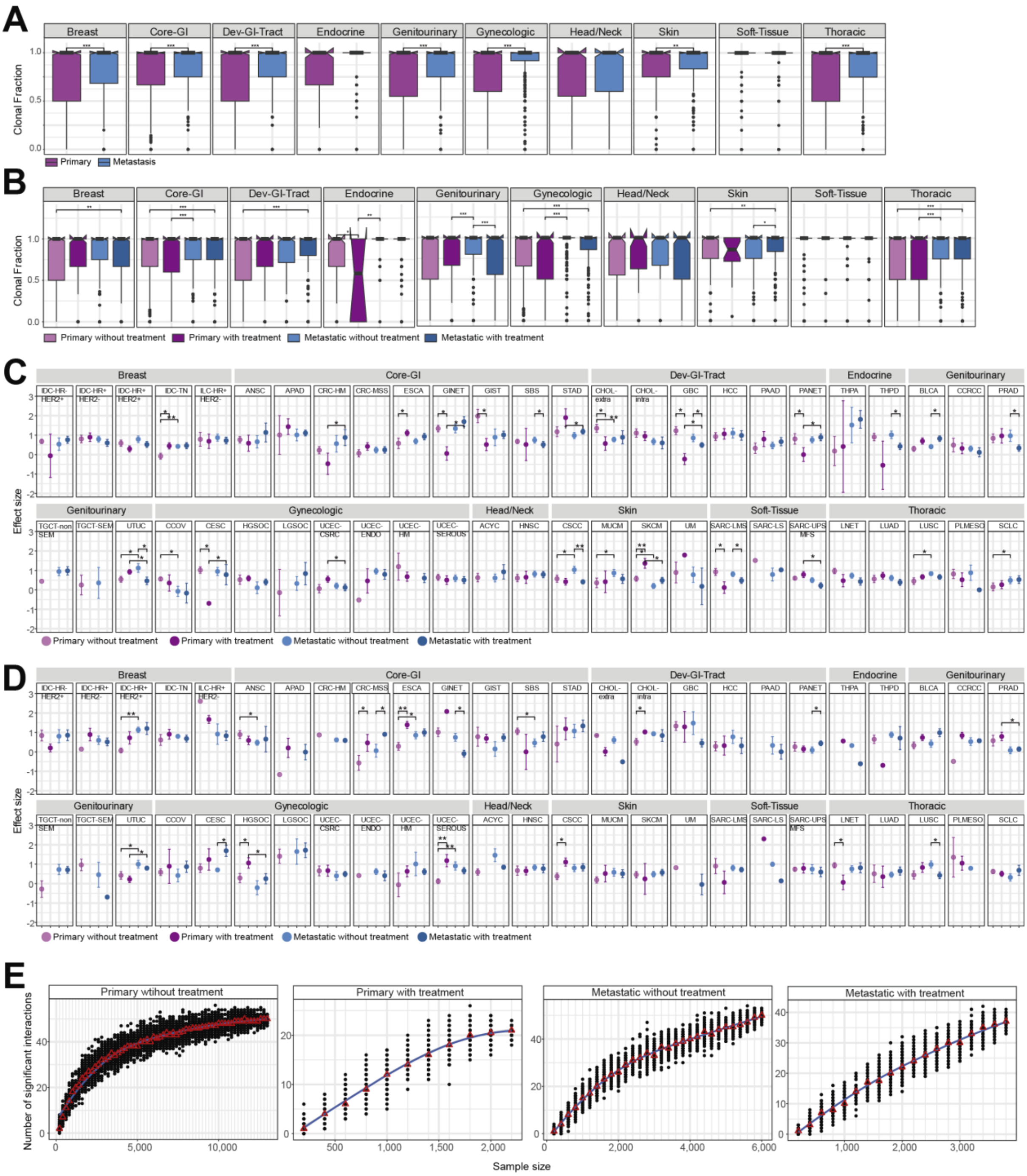


**Figure S6**. Tumor clonality distribution according to the (**A**) sample status and (**B**) treatment conditions across cancer types. In boxplots, the centre line represents the median, the box limits indicate the first and third quartiles, and the whiskers extended to the lowest or highest data points within 1.5 times the interquartile range from the quartile. *P*-value are calculated using Mann-Whitney *U* test, with asterisks denoting statistical significance (∗ *P*-value < 0.05). (**C**-**D**) The average interaction strength between mutations and CNA within a gene in the CNA-loss and CNA-gain models is depicted depending on the treatment conditions at the subtype level. Error bars indicate the standard error within each condition. Statistically significant differences among conditions were determined using the Mann-Whitney *U* test. **(E)** Saturation analysis by subsampling both primary and metastatic tumors, divided based on treatment conditions. Each data point represents the number of significantly detected interaction by randomly selected subsamples, and red triangle denotes the median number of significant two-way interactions at FDR 10% as the sample size increases across multiple runs (*N*=200, 100 runs). Smoothing lines (blue) were added using the 'loess' smoothing method to visualize the number of significant interactions and the red triangle represents the median number of significant interactions.

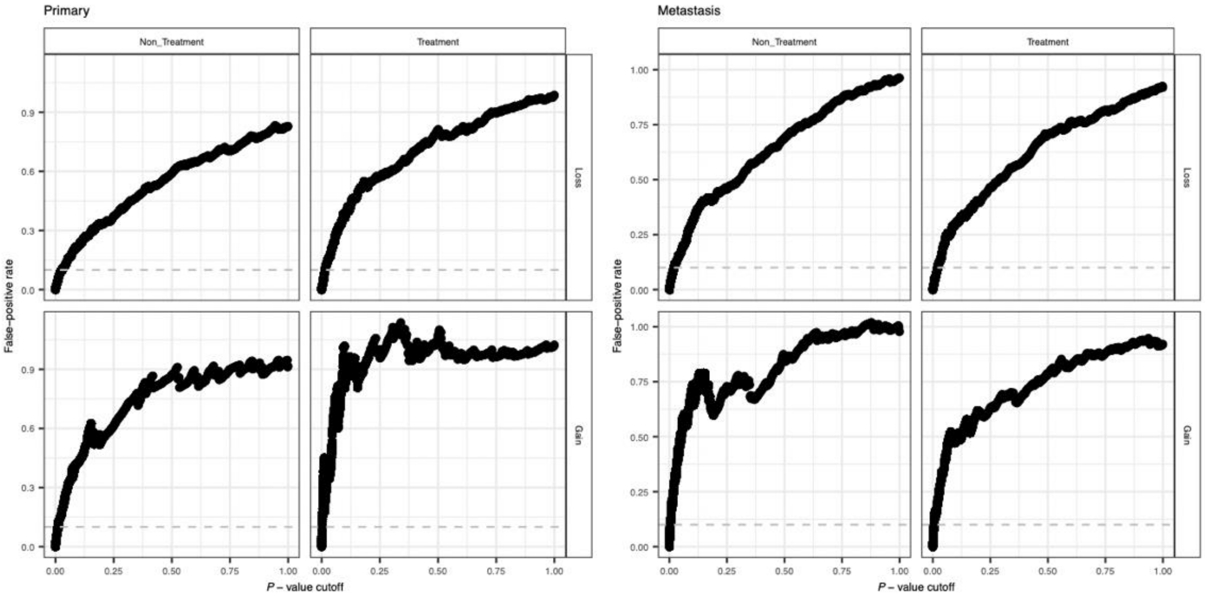


**Figure S7**. False-positive discovery rate (FDR) assessment for two-way interactions based on treatment status. The FDR values were calculated at various *P*-value thresholds using a permutation approach involving 100 iterations. This permutation strategy was employed to assess the significance of co-occurrence between pairs of genomic alterations within the same gene, while controlling for genomic heterogeneity within and across samples and genes. The strategy maintained the total number of alterations for each genomic event across samples as well as the total number of alterations per sample. Each genomic event (mutation, CNA-loss, and CNA-gain) was treated as a separate class and analyzed independently within each cancer type and state. For each specified *P*-value cut-off, the FDR was estimated as the ratio between the number of detected interactions in the permuted data matrix (false-positive interactions) and the number of interactions detected in the original data matrix (true observed interactions). The resulting average FDR values are plotted for each *P*-value cut-off for primary (left panel) and metastatic models (right panel).

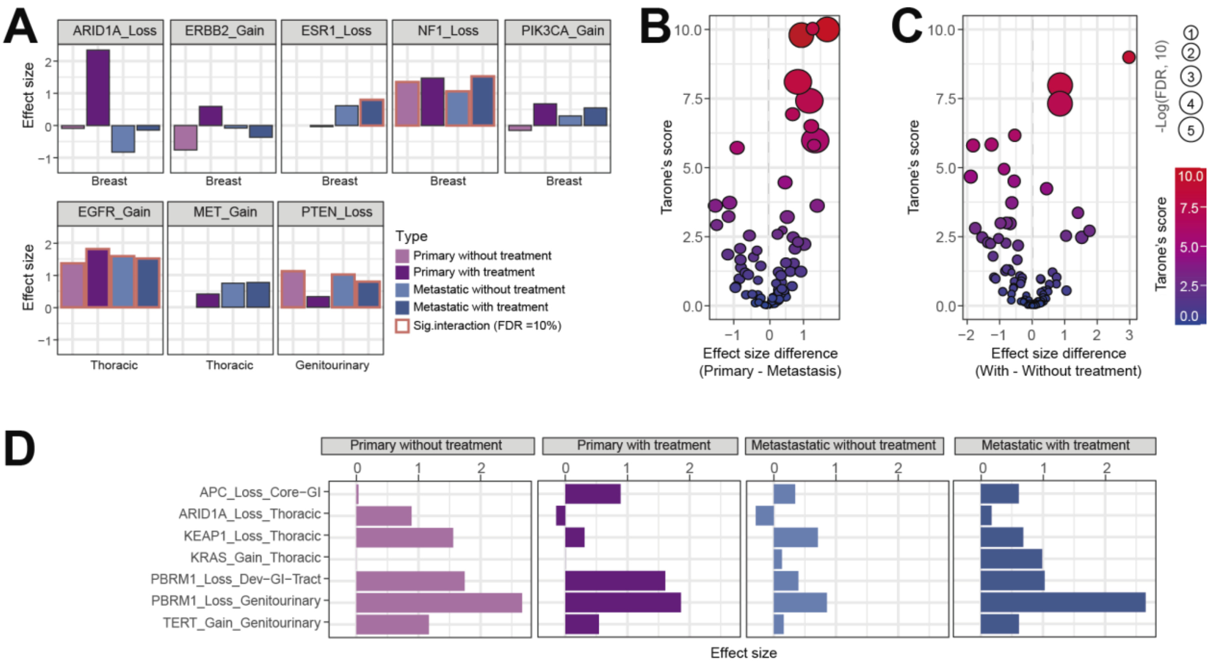


**Figure S8**. Effects of treatment conditions on the interaction strength of co-occurrences between mutations and CNA within a gene across cancer states. (**A**) Distribution of interaction strengths for known drug-resistant genes in their targeted cancer types under different treatment conditions and cancer states. (**B**-**C**) Condition-specific interaction strength distribution. 97 pairs in treated versus non-treated tumours and 84 pairs in primary versus metastatic tumours were tested using the odds ratio heterogeneity test (Tarone's test). Volcano plot comparing differences in interaction strengths between two conditions. (**D**) Interaction strength between mutations and CNA within a gene was assessed for seven significantly different interactions detected by the Tarone's test (FDR at 10%).

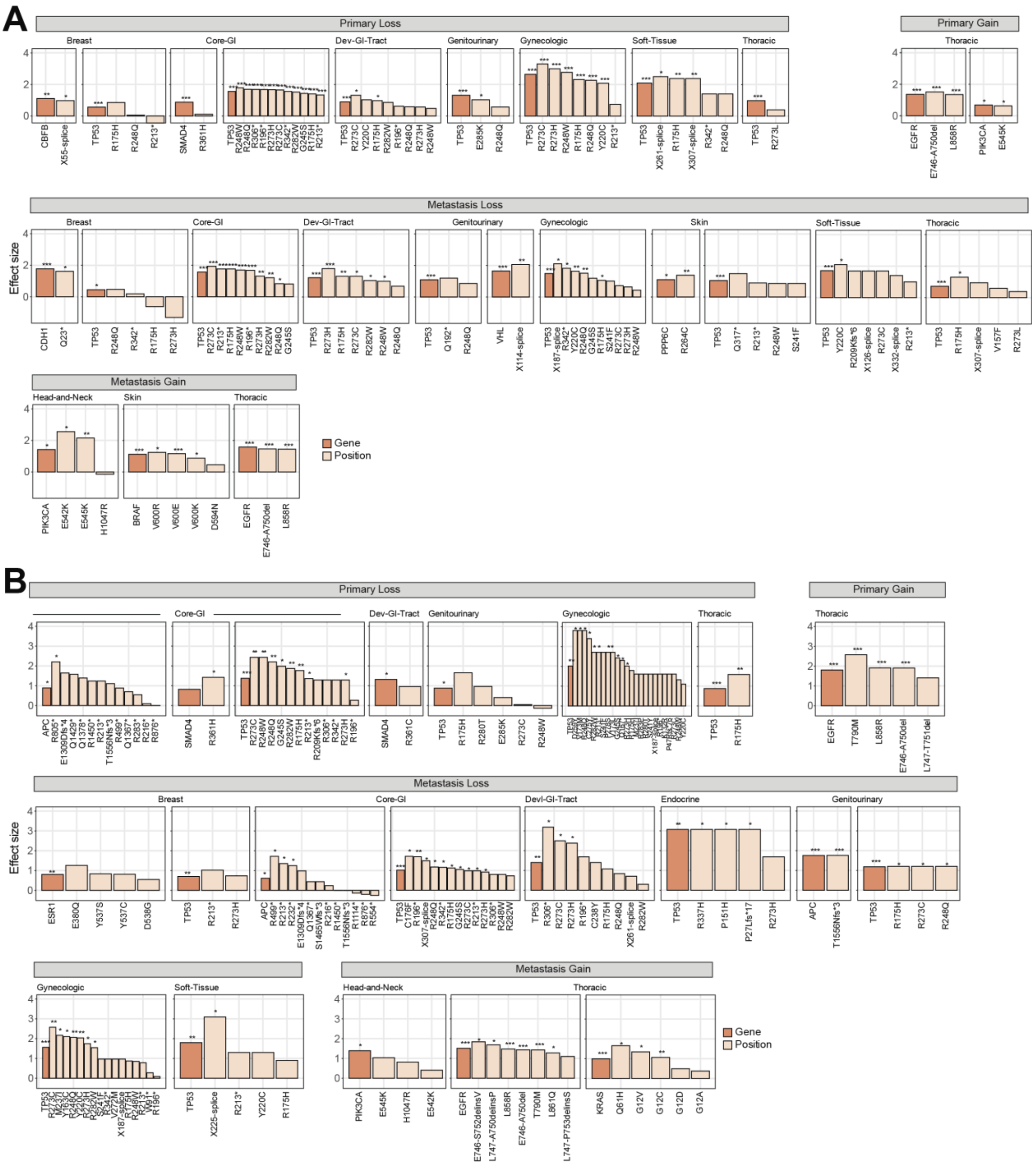


**Figure S9**. Position-specific interaction strength between mutation and CNAs within a gene across cancer types. Interaction strength distribution of detected two-way interactions at FDR 10% in (**A**) treated and (**B**) non-treated conditions at highly mutated positions (mutation frequency > 1% within the detected cancer type).

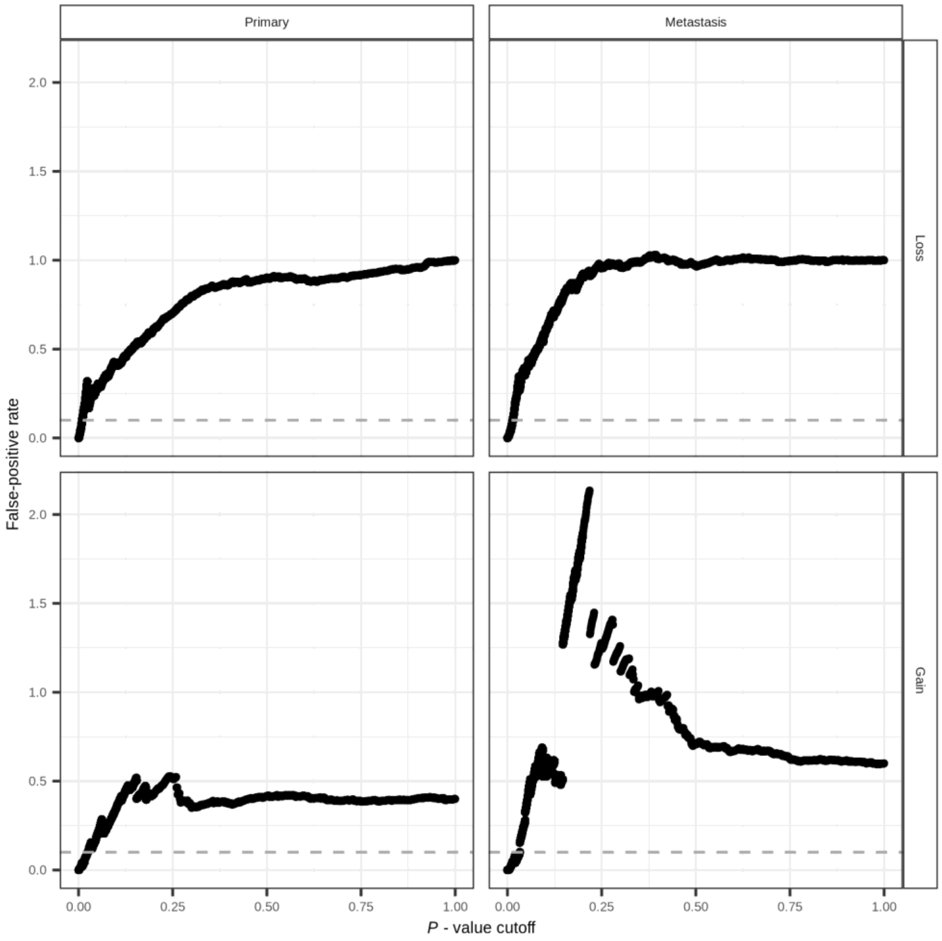


**Figure S10**. False-positive discovery rate (FDR) assessment for three-way Interactions in primary and metastatic tumors. The FDR values were calculated at various *P*-value thresholds using a permutation approach involving 100 iterations. This permutation strategy was employed to assess the significance of the co-occurrence of a pair of genomic alterations in the first gene depending on the second gene (three-way interaction), while controlling for genomic heterogeneity within and across samples and genes. The strategy maintained the total number of alterations for each genomic event across samples as well as the total number of alterations per sample. Each genomic event (mutation, CNA-loss, and CNA-gain) was treated as a separate class and analyzed independently within each cancer type and state. For each specified *P*-value cut-off, the FDR was estimated as the ratio between the number of detected interactions in the permuted data matrix (false-positive interactions) and the number of interactions detected in the original data matrix (true observed interactions). The resulting average FDR values are plotted for each *P*-value cut-off for primary (left panel) and metastatic models (right panel).

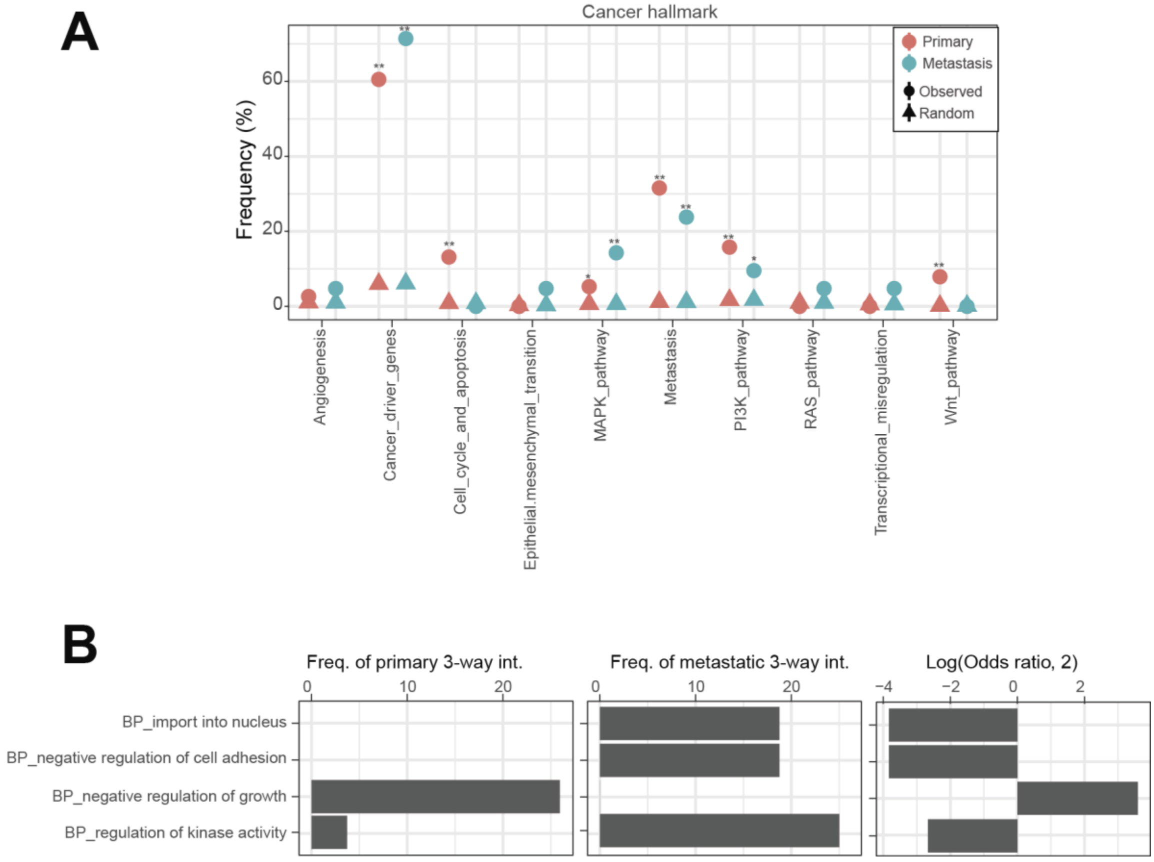


**Figure S11**. Functional annotation of three-way interactions (**A**) The overlapping frequency of three-way interaction pairs in primary and metastatic tumors was analyzed with respect to cancer hallmarks (Li et al., 2023). For the random set, two cancer genes from the MSK-panel were randomly selected, matching the number of tested pairs (38 random pairs in primary and 21 random pairs in metastatic tumors). A total of 10,000 randomizations were performed, and the empirical *P*-value was determined as the proportion of randomizations where the detected overlapping frequencies exceeded those observed in the real data. (**B**) Gene Ontology enrichment analysis was conducted for three-way interactions specifically observed in primary and metastatic tumors (Fisher's one-tailed test with a significance level of $P < 0.05$). The enrichment preference for functional annotations was calculated as the odds ratio of primary versus metastatic tumors, with a higher odds ratio indicating stronger enrichment in metastatic tumors compared to primary tumors.

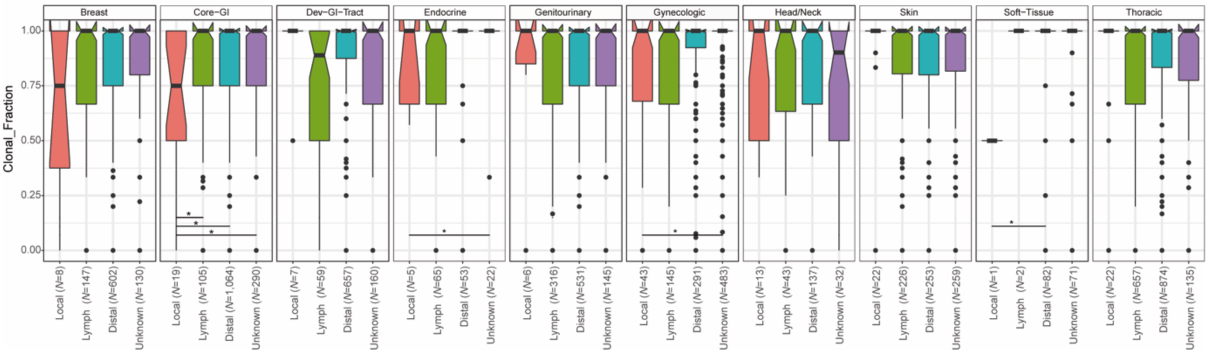


**Figure S12**. Distribution of tumour clonality by metastatic biopsy location across cancer types. In each boxplot, the centre line represents the median value, while the box limits indicate the first and third quartiles. The whiskers extend to the lowest and highest data points within 1.5 times the interquartile range from the first quartile. The *N* value indicates the number of samples included in the analysis. The *P*-value corresponds to the Mann–Whitney *U* test, with asterisks denoting statistical significance (**P*-value < 0.05).
